# An integrated biomaterials-centred approach of the ageing thymic microenvironment reveals design principles for regenerative biomaterials

**DOI:** 10.64898/2026.07.27.740917

**Authors:** A. Di Bernardo, C. Zanirato, F. Briatico, P. Petrini, C. S. Butnarasu, F. Leo, C. Ambrogio, E. Patrucco, A. Locatelli, F. Oliva, A. Passoni, C. Medana, S. Visentin, L. Sardelli

**Affiliations:** Department of Molecular Biotechnology and Health Science, University of Torino, Torino, Italy; Department of Chemistry, Materials, and Chemical Engineering “G. Natta”, Politecnico di Milano, Milan, Italy; BioAvatar Lab, Department of Chemistry, Materials, and Chemical Engineering “G. Natta”, Politecnico di Milano, Milan, Italy; Department of Oncology, University of Turin, San Luigi Gonzaga University Hospital, Orbassano, Italy; Thoracic Surgery Division, Department of Oncology, University of Turin, San Luigi Gonzaga University Hospital, Italy; Istituto di Ricerche Farmacologiche Mario Negri IRCCS, Milan, Italy

**Keywords:** Immune system, ageing, mechanical properties, rheology, lipidome

## Abstract

Thymic involution is commonly addressed as a loss of epithelial and lymphoid tissue, yet the accompanying remodelling of the microenvironment remains poorly defined. This work applied a biomaterials-centered approach to compare young and aged bovine thymus by integrating histology, oscillatory rheology, untargeted lipidomics, ICP–MS and AP-MALDI mass spectrometry imaging. This holistic approach connects the mechanical, compositional, and spatial features of the native thymus with the development of thymus-inspired biomaterials. Ageing increased both storage and loss moduli by more than one order of magnitude and reduced the linear viscoelastic region approximately fivefold, defining a markedly stiffer and more strain-sensitive material state. This mechanical transition was accompanied by lipid remodelling, with double realtive contribution of triacylglycerols to the lipid pool doubled and loss of membrane-associated phospholipids. The elemental profile also contracted, with total metal content decreasing by one-third and zinc showing a reduction of 70%. At the architectural level, the corticomedullary ratio was more than halved, while AP-MALDI imaging revealed an approximately 40% reduction in the annotated molecular repertoire and a shift from homogeneous to fragmented distributions of choline and representative phosphatidylcholine species. These results show that thymic ageing emerges from the concomitant variations of viscoelastic behaviour, chemical composition, and molecular organization. By defining these interconnected alterations, this study establishes a materials-based foundation for the design of bioinspired systems aimed at reproducing features of the young thymic niche and supporting future thymic repair.

**Highlights:**

- Ageing increased viscoelastic moduli and reduced deformation tolerance.
- The lipid profile shifted from membrane lipids towards triacylglycerols.
- The elemental pool contracted, with zinc showing the strongest depletion.
- Histology and AP-MALDI-MSI revealed loss of spatial organization.

**Graphical abstract:** 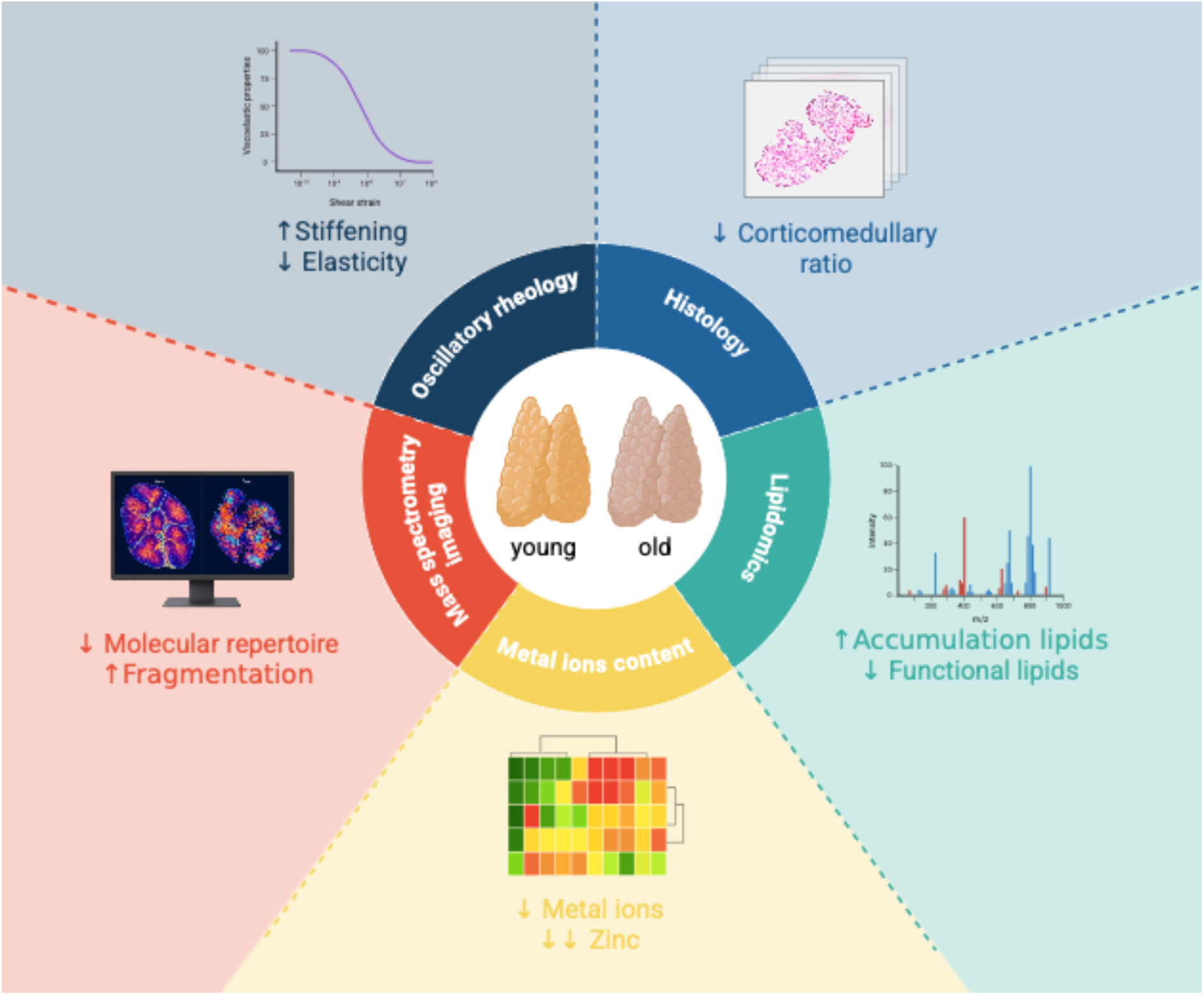

## 1. Introduction

Population ageing is reshaping medicine[1]. By 2050, people aged 60 years or older will account for 22% of the global population[2], [3]. Yet increased longevity remains accompanied by chronic disease and diminished resilience to infection, cancer and systemic injury[4]. In this context, the human thymus has a central role as the primary site of T-cell maturation, generating the diverse, self-tolerant repertoire of naïve T cells required for adaptive immunity[5], [6]. However, the thymus is also among the organs most profoundly affected by age-associated involution, progressively losing functional epithelial space and ultimately undergoing extensive adipose replacement[4]. For decades, this rapid involution supported the view that the thymus was largely dispensable in adulthood and a developmental organ whose essential function had been completed early in life. This view is now changing[7]. In a 2026 study of more than 27,000 adults, poorer radiographic thymic health was associated with increased all-cause and cardiovascular mortality and a higher incidence of lung cancer. A separate study of approximately 3,500 patients receiving immune-checkpoint inhibitors linked thymic health to treatment outcomes across several cancer types[8], [9]. These findings position the thymus as a compelling target for regenerative medicine and have attracted growing interest from both public funding bodies and private biotechnology companies. Engineering thymic tissue by restoring a youthful microenvironment could strengthen immune competence against infection and cancer, positioning this often-overlooked organ as a key contributor to systemic health and healthy longevity[4], [7], [10], [11].

Current thymic regenerative strategies have predominantly focused on cellular and soluble-factor-based interventions, including cytokine and endocrine modulation (e.g., IL-7, IL-22, growth-hormone-associated approaches and sex-steroid blockade), as well as precursor-cell therapies, pluripotent-stem-cell-derived thymic epithelial progenitors, FOXN1 reprogramming and organoids[12], [13], [14], [15]. By contrast, scaffold-assisted tissue-engineering approaches have largely been limited to decellularized extracellular-matrix scaffolds and GelMA-based hydrogels. Nevertheless, these studies have demonstrated that engineered three-dimensional matrices can enhance thymopoiesis and support thymic-cell survival[16], [17]. Despite these advances, the thymic microenvironment remains largely a biological black box and durable thymic regeneration, however, remains elusive[18], [19]. Consequently, it is still unclear which physical and compositional features of the thymic microenvironment should be preserved, targeted or restored through biomaterial design to support thymic regeneration[20]. Individual studies have begun to open this black box, but have provided only fragmented observations. For example, although ageing-associated extracellular-matrix remodelling is known to alter tissue stiffness and viscoelasticity in several organs[21], whether comparable changes occur in the aging thymus has not been investigated. This represents an important gap from a biomaterials perspective, because both resident and infiltrating thymic cells respond to mechanical cues that influence their behaviour and function.

Mechanical characterization alone, however, cannot fully define the thymic microenvironment, because the regenerative properties of a material also depend on its biochemical composition. At the macroscopic level, thymic involution is accompanied by progressive adipose replacement[22], [23]. At the molecular level, targeted lipidomics studies have identified age-associated changes in selected lipid species[24], [25]. However, an untargeted characterization of the ageing thymic lipidome is still lacking, leaving the full extent of lipid remodelling unresolved.

The thymic elemental composition has received even less attention. Ageing disrupts metal-ion homeostasis across multiple tissues, with consequences for redox balance, mitochondrial metabolism, enzymatic activity and cellular signalling[26], [27], [28], [29]. In the thymus, zinc supports thymulin activity, T-cell differentiation and thymic regeneration[30], [31], [32]. However, age-associated changes in the broader elemental landscape of the organ have yet to be comprehensively assessed. It therefore remains unclear which lipid and elemental features represent meaningful targets for the design of thymus-inspired biomaterials.

Because thymic function depends on spatially compartmentalized cortical and medullary niches, biomaterial design requires an understanding not only of which molecular cues are present, but also of where and how they are presented within the tissue. In this view, MALDI mass-spectrometry imaging has demonstrated that molecular distributions can be spatially resolved within thymic tissue[33], while metabolic changes have also been detected in the ageing murine thymus[34]. Yet most thymic metabolomics and imaging studies have examined pathological or experimentally perturbed conditions[35], [36], [37], and the spatial molecular reorganization accompanying physiological ageing has not been systematically characterized. Hence, thymic ageing extends beyond epithelial loss and adipose replacement to encompass broader physicochemical remodelling of the tissue. Defining age-associated changes in the mechanical, compositional and spatial properties of the thymic microenvironment could identify the physicochemical parameters to guide the design of biomaterials capable of recreating a young regenerative niche[38], [39].

To address these gaps, we used calf and adult cow thymic tissue as a large-animal comparative model of age-associated thymic involution. This comparison has previously been employed to investigate age-dependent molecular changes in bovine thymus, while studies across cattle age groups have documented the progressive morphostructural involution of the organ [40], [41]. The availability of large tissue volumes further enables complementary mechanical, compositional and spatial analyses within the same experimental framework. By providing an integrated characterization of young and aged thymic tissue using histology, oscillatory rheology, untargeted lipidomics, elemental profiling and high-resolution mass-spectrometry imaging (**Fig.1**), this work aims to establish a holistic, microenvironment-centred approach for understanding thymic ageing and hence supporting the design of thymus-inspired regenerative biomaterials.

**Fig. 1.**
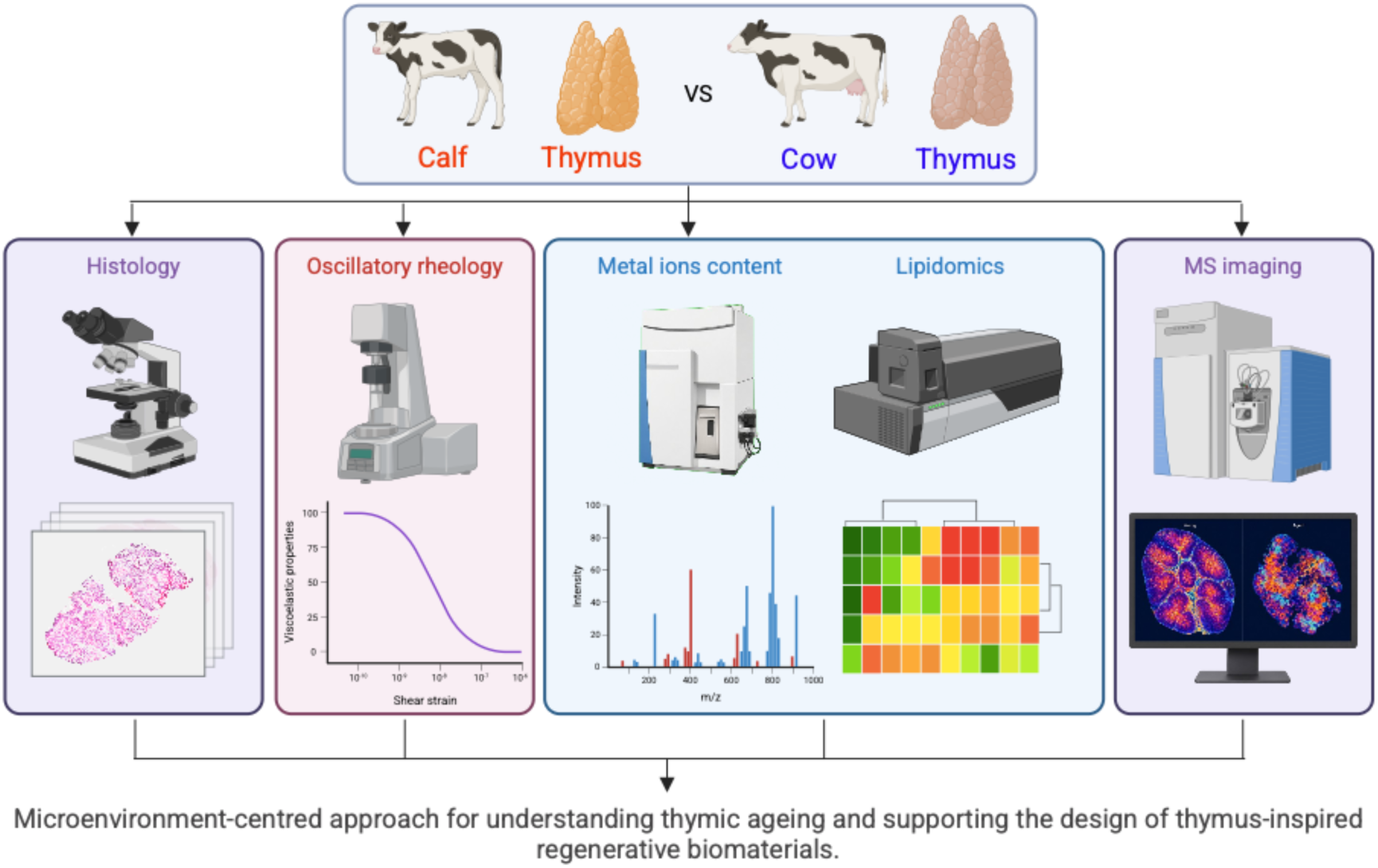
schematic representation of the work rationale

## 2. Materials and methods

### 2.1 Materials

Thymi from calves (7-9 months old) and cows (20-24 months old) were obtained from a local abattoir and preserved in 2% (w/v) sodium citrate solution to prevent coagulation during transport and handling.

Methanol, acetonitrile, 2-propanol, chloroform, ammonium formate, and formic acid were of LC–MS or analytical grade and were purchased from Sigma-Aldrich/Merck (Milan, Italy). The EquiSPLASH™ Lipidomix mixture of deuterated lipid internal standards was purchased from Avanti Polar Lipids (Alabaster, AL, USA) and used for class-matched normalization of lipid peak areas. Nitric acid (70%, purified by reditillation, trace-metal grade) was obtained from Sigma-Aldrich/Merck (Milan, Italy). Multi-element standard solutions for ICP-MS calibration were purchased from Areachem (Melito di Napoli, Napoli, Italy). α-Cyano-4-hydroxycinnamic acid and trifluoroacetic acid were purchased from Merck LifeScience S.r.l (Milan, Italy). Ultrapure water was produced using a Milli-Q water purification system (Merck Millipore, Darmstadt, Germany).

### 2.5 Histology: Hematoxylin and Eosin Staining

Fresh thymic tissue from calves and cows was collected and stored at -80°C until sectioning. Cryosections of 10 μm thickness were cut at -20 °C using a cryostat (Leica CM1860 UV, Leica Biosystems) and mounted onto slides, which were stored at -80°C until staining. Before staining, sections were post-fixed in 4% paraformaldehyde for ten minutes and rinsed in distilled water. Sections were stained with Mayer’s hematoxylin for 4 minutes, rinsed in water, and counterstained with eosin for 1 minute. Sections were subsequently dehydrated through a graded ethanol series, cleared in xylene, and mounted with Eukitt mounting medium before coverslipping. Stained sections were examined, and images were captured with the Virtual Slide Microscope (Olympus).

### 2.2. Rheological characterization

Tissue samples were cut into discs of approximately 25 mm in diameter using a biopsy punch. Samples were either analyzed immediately or, upon sodium citrate solution removal, were frozen at80 °C and stored for different time periods (*i.e.*, 1 week and 3 weeks). Before measurements, frozen samples were gently thawed at4 °C.

Mechanical characterization was performed using a modular rheometer (MCR 502e, Anton-Paar, AT) equipped with a 25 mm parallel-plate geometry with serrated plates to prevent sample slippage. All measurements were carried out at 25°C. Prior to rheological measurements, a normal pre-load of 2 N was applied for 3 min to ensure proper contact between the sample and the plates and to allow stress relaxation and then maintained during the measurement duration. Time sweep tests were carried out by applying different normal forces, from 2 to 8 N (corresponding to 1 to 4 kPa of normal stress), as pre-loads for 3 min, followed by oscillatory shear at 0.04% strain amplitude and 1 rad·s^-^¹ for 5 min.

Rheological characterization started with an amplitude sweep test at a constant oscillation frequency of 1 Hz, with shear strain ranging from 0.001% to 10%, in order to determine the linear viscoelastic region (LVR). Then, frequency sweep tests were performed within the LVR (γ = 0.04%) by varying the angular frequency from 1 to 100 rad·s^-^¹.

### 2.3 Metal ions content via Inductively Coupled Plasma-Mass Spectrometry

200 mg of thymic samples were weighed and placed into a 10 mL acid-washed graduated polypropylene tube. Subsequently, 1 mL of 70% (v/v) nitric acid was added. The samples were left semi-capped under a fume hood to initiate acid hydrolysis for 48 hours at room temperature. Then, the sample was transferred into a microwave digestion vessel. A microwave-digestion system (Ethos UP, Milestone, Bergamo, Italy) was used for mineralization according to an optimized protocol. Specifically, the heating ramp involved a 10 min increase to 180 °C, followed by a 20-min isothermal period at 180 °C, after which the system was allowed to cool to room temperature. After digestion, the mineralized sample was transferred into a 10 mL graduated polypropylene tube. The digestion vessel was rinsed with 3 mL of ultrapure water, and the rinse was combined with the mineralized sample. The resulting solutions were filtered through 0.45 μm hydrophilic PTFE syringe filters to remove insoluble residue and diluted with ultrapure water to obtain a final nitric acid concentration of 4% (v/v). Quantification of elemental concentrations was performed using calibration curves from certified reference standards. Elemental analysis was carried out using an iCAP-Q ICP–MS system (Thermo Fisher Scientific) operating in standard plasma conditions (RF power 1.5 kW) with a cooled quartz cyclonic spray chamber, nickel sample and skimmer cones. To minimize spectral interferences arising from polyatomic species, measurements were conducted in helium collision mode with a gas flow of 5 mL/min and optimized bias voltages (−18 V pole bias and−21 V CCT bias). Following instrument tuning and calibration, metal concentrations in thymic samples were determined directly from the established calibration curves.

### 2.4 Untargeted lipidomics

Thymic tissue samples (approximately 10 mg) were weighed and subjected to initial homogenization in ice-cold methanol at a solvent-to-tissue ratio of 20 µL/mg. Homogenization was achieved by vortexing the mixture for 30 s, followed by sonication for 5 min and a subsequent incubation period of 5 min at 4°C to ensure complete disruption of cellular structures and efficient lipid extraction. The homogenates were then centrifuged at 16,000 × g for 5 min at 4°C, and 100 µL of the clear supernatant was transferred to a new microcentrifuge tube. Subsequently, a modified Bligh and Dyer extraction protocol was employed[42]. To the supernatant, 300 µL of methanol, pre-spiked with a mixture of deuterated lipid internal standards to a final concentration of 250 ppb (EquiSPLASHTM, Avanti Research), was added, followed by 400 µL of chloroform and 320 µL of ultrapure water. The mixture was vortexed vigorously for 30 s to promote phase separation, then centrifuged again at 16,000 × g for 5 min at 4°C. The lower organic phase, containing the extracted lipids, was collected and evaporated to dryness under nitrogen flow at room temperature. The dried lipid extracts were finally reconstituted in 200 µL of solvent B (as defined in the chromatographic method), yielding samples ready for untargeted lipidomics analysis by liquid chromatography-high resolution mass spectrometry (LC-HRMS). Lipid profiling was performed using an LC–Orbitrap MS platform consisting of a Vanquish Core HPLC system coupled to an Orbitrap Exploris 120 mass spectrometer (Thermo Fisher Scientific, Waltham, MA, USA) equipped with a heated electrospray ionization probe (H-ESI). Chromatographic separation was achieved on a Luna C18 column (150 × 2.0 mm, 3 μm particle size, 100 Å pore size; Phenomenex, Torrance, CA, USA) maintained at 50 °C, with a flow rate of 0.25 mL/min. The sample injection volume was 20 μL. The mobile phase consisted of solvent A (H₂O:acetonitrile, 40:60 v/v, containing 1 mM ammonium formate and 0.1% formic acid) and solvent B (isopropanol:acetonitrile, 90:10 v/v, containing 0.1% formic acid). Lipids were separated using a 40 min gradient: 40% B from 0– 2 min, increased to 100% B from 2–35 min, held at 100% B from 35–40 min, followed by an 8 min re-equilibration at 40% B. After separation, mass spectrometric analysis was performed using an electrospray ionization (ESI) source operating in positive ion mode with a spray voltage of +3.5 kV and ion transfer tube temperature of 300 °C. Full-scan MS spectra were acquired at a resolution of 60,000 over an *m/z* range of 150–1100. Data-dependent acquisition (DDA) was employed for MS/MS fragmentation using higher-energy collisional dissociation (HCD) with a collision energy of 28 a.u.. Raw data were processed using Xcalibur (Thermo Fisher Scientific) and MS-DIAL version 4.9 for peak detection, alignment, and lipid annotation against the LipidBlast MSP spectral library. Lipid annotations were reported at the lipid species sum-composition level. Peak area integration was subsequently performed using Skyline software. Peak areas were normalized to the corresponding class-matched deuterated internal standard included in the EquiSPLASH mixture, and the resulting normalized areas were used for relative lipid abundance analysis.

### 2.6. AP-MALDI-MSI Analysis

Cryosections of 10 μm thickness were cut from thymus tissue of calves (n=3) and cows (n=3) at -20°C using a cryostat (Leica CM1860 UV, Leica Biosystems) and mounted onto AP-MALDI target plates. α-cyano-4-hydroxycinnamic acid (CHCA) matrix (10 mg/mL in 70% ACN, 0.1% TFA) was deposited using a SunCollect MALDI sprayer (SunChrom GmbH, Friedrichsdorf, Germany) with the following parameters: a spray speed of 800 mm/min, line distance of 2 mm, 20 layers, and a flow rate of 80 μL/min, resulting in a final concentration of 10 μg/mm². Samples were dried under vacuum prior to MALDI analysis. MSI was performed using an AP-MALDI ion source (MassTech Inc., Maryland, USA) coupled to an Orbitrap Q-Exactive mass spectrometer (Thermo Scientific, Bremen, Germany). Experiments were conducted using Target-ng software (MassTech Inc.) with the source operating in continuous motion mode. For each experimental group, one replicate was acquired at 50 μm spatial resolution and the others at 150 µm. The mass spectrometer was operated in Full MS positive ionization mode over an m/z range of 60–800, at a resolving power of 70,000 (at m/z 200). Automatic gain control (AGC) was set to 5 × 10⁶ with a maximum injection time of 500 ms. The capillary temperature was maintained at 250 °C. Raw MALDI-MSI data were converted to imzML format using MT imzML Converter (light) 0.0.8. For metabolite identification, imzML data were uploaded to METASPACE Pro, and metabolite annotation was performed using four databases: HMDB-v4, CoreMetabolome-v3, KEGG-v1, and LipidMaps-2017-12-12, applying a false discovery rate (FDR) threshold of 20%. For each experimental group, only metabolites detected in at least two of the three biological replicates were retained.

### 2.7 Statistical analysis

Experiments were performed at least in biological triplicate, and data were reported as mean ± standard deviation unless otherwise specified. The normality of data distribution was assessed using the D’Agostino–Pearson test. Depending on the distribution, comparisons between two groups were performed using either Student *t*-test or Mann–Whitney test, while comparisons among multiple groups were performed using one-way ANOVA or the Kruskal–Wallis test. Statistical analyses were carried out using GraphPad Prism 9 (GraphPad Software, USA). Differences are shown in the graphs only when statistically significant. Statistical significance was defined as *p* < 0.05 (\**), p < 0.01 (**), p < 0.001 (\*\*\**), and *p < 0.0001 (****)*.

## 3. Results

### 3.1 Architectural variation of the thymic microenvironment

Histological analysis revealed distinct differences in thymic architecture between calves and cows. Calf thymus displayed a compact and well-preserved lobular organization, with a prominent, densely stained cortical compartment surrounding smaller and lighter medullary regions (**Fig. 2a**). In cow thymus, the parenchyma appeared more discontinuous, with a lower relative predominance of the cortical compartment and less evident corticomedullary compartmentalization (**Fig. 2b**). Consistently, the cortex-to-medulla area ratio decreased from 10.3 in calves to 4.7 in cows (**Fig. 2c**). Comparable histological features were observed across the three animals analysed.

**Fig. 2.**
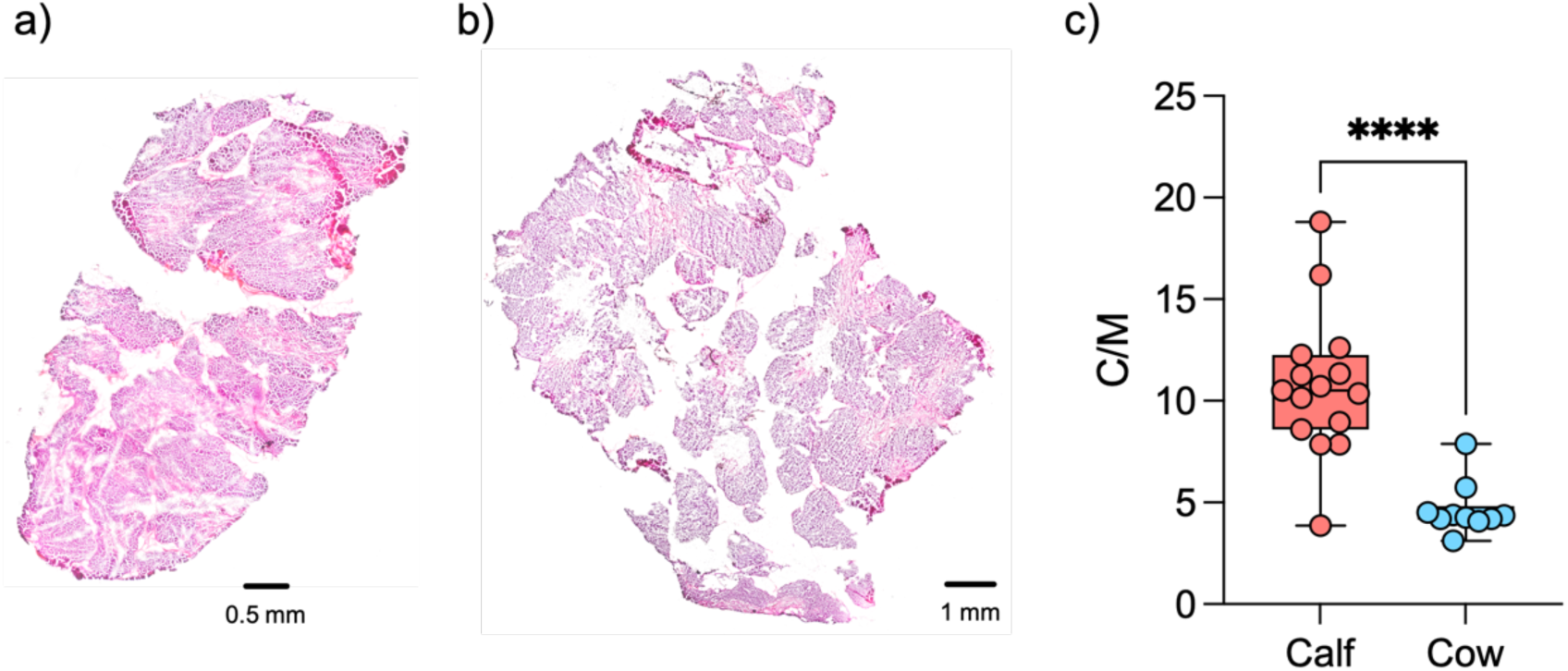
Representative histological images of calf (a) and cow (b) thymus used to quantify the cortico-medullary ration (c).

### 3.2 Age-associated stiffening and reduced deformation tolerance define distinct thymic material states

Thymic biomechanics remains largely uncharacterized, particularly in relation to ageing. As a first step towards a biomaterials-centred investigation of thymic involution, the viscoelastic behaviour and deformation tolerance of calf and cow thymic tissues were therefore compared. Independently of age, fresh thymus tissues exhibited a solid-*like* behaviour, with G’ >> G’’ across the entire range of shear strain amplitudes considered (**Fig. 3b**). Aging was associated with the linear viscoelastic region (LVR), with calf samples exhibiting a more extended LVR, reaching a limiting shear strain amplitude of 0.2%, compared to cows that were limited to 0.04%. For this reason, a shear amplitude of 0.04% was selected for all subsequent rheological characterizations.

**Figure 3.**
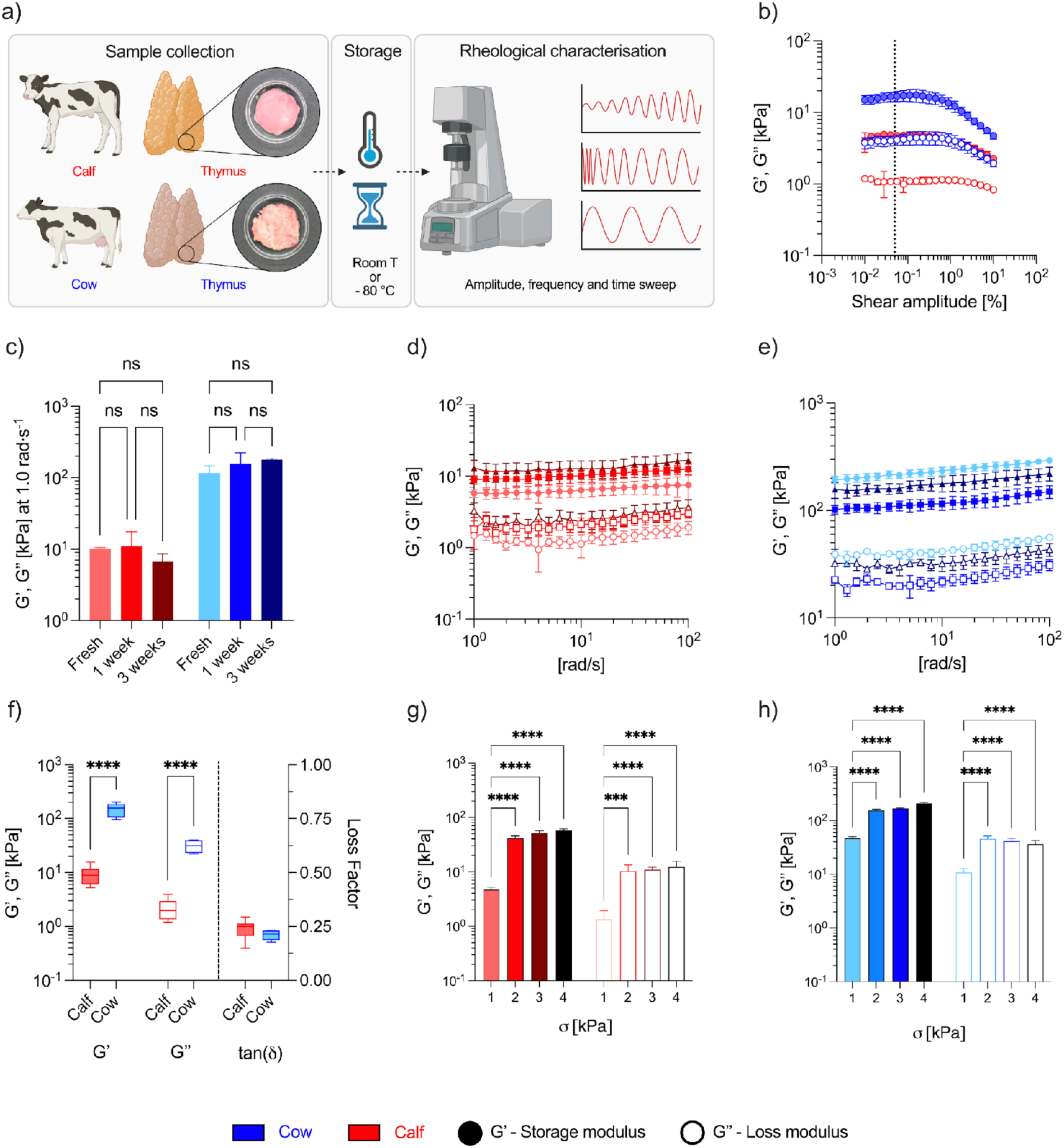
(**a**) Schematic overview of the rheological characterization performed on thymus samples obtained from calves (7–9 months old) and cows (20-24 months old). (**b**) Amplitude sweeps tests performed to identify the linear viscoelastic region (LVR). (**c**) Storage modulus (G′) of calf (red) and cow (blue) thymus under different storage conditions. (**d, e**) Frequency sweep profiles of three calf and cow thymus samples, respectively. Error bars indicate the variation among technical replicates obtained from thymus of the same animal. (**f**) Storage modulus (G′), loss modulus (G″), and tan δ extracted at 1 rad·s^-^¹. (**g, h**) Storage and loss moduli of calf and cow thymus, respectively, measured at the final time point of time sweep tests performed under different applied normal stresses.

The effect of storage was evaluated by analysing the G’ and G’’ trends in the angular frequency spectra over the range of 1–100 rad·s^-^¹ (**Fig. S1**) and by comparing the values at 1 rad·s^-^¹ (**Fig. 3c**). Under all storage conditions, thymic tissues maintained a predominantly elastic response, with G′ remaining higher than G″. Compared with fresh samples, storage at −80 °C for 1 or 3 weeks was associated with a slight decrease in G′ and G″ in calf thymus and an opposite trend in cow thymus; however, these changes were not statistically significant, and no effect of storage duration was observed. In calf thymus, G′ remained essentially stable after 1 week of storage, varying by less than 10% compared with fresh tissue values. After 3 weeks, G’ remained within the same order of magnitude as fresh tissue, ranging from 10.08 ± 0.48 kPa in fresh samples to 6.66 ± 1.99 kPa after 3 weeks. A similar behaviour was observed for G″, which showed limited variation across storage conditions, ranging from 2.52 ± 0.16 kPa in fresh samples to 1.37 ± 0.24 kPa after 3 weeks. Similarly, in cow thymus, freezing did not significantly affect the viscoelastic moduli, with both G′ and G″ remaining within the same order of magnitude across all storage conditions. Importantly, the predominantly elastic behaviour of the tissue was preserved in all conditions, with G′ consistently higher than G″ throughout the analysed frequency range (**Fig. S2**).

Inter-animal and intra-animal variability were assessed by measuring three thymic lobules from each of three independent animals and by comparing the corresponding G′ and G″ spectra over the frequency range of 1–100 rad·s^-^¹ (**Fig.3 d-e; Fig.SI 2**). For both calf and cow thymus, all samples maintained a predominantly elastic behaviour, with G′ consistently higher than G″ across the analysed frequency range. In calf thymus, inter-animal differences accounted for 67% of the variability in G′ and 56% of the variability in G″, while intra-animal variability accounted for 33% and 44%, respectively. In cow thymus, the inter-animal component was even more pronounced for G′, accounting for 85% of the variability, with only 15% attributable to intra-animal differences. By contrast, G″ variability in cow thymus was almost equally distributed between inter-animal and intra-animal components, which accounted for 52% and 48%, respectively. Thus, variability in G′ was mainly associated with differences between animals, particularly in cow thymus, while variability in G″ showed a more balanced contribution of inter-and intra-animal components.

The effect of ageing was quantified at a fixed angular frequency of 1 rad s⁻¹ to enable direct comparison between groups (**Fig. 3f**), although the same trend was observed across the entire frequency range tested. Cow thymus exhibited markedly higher values for both viscoelastic components than calf thymus, with G′ increasing approximately 16.4-fold, from 9 to 151 kPa, and G″ increasing 14.4-fold, from 2 to 31 kPa. Thus, ageing was associated with a more than one-order-of-magnitude increase in both the elastic and viscous components of thymic tissue. Despite this pronounced stiffening, tan(δ) comparable between calf and cow thymus, with mean values around 0.2, indicating that the predominantly elastic nature of the tissue was maintained, and that the relative importance of conservative and dissipative contribution to mechanical response were unaffected by structural and compositional variation associated to aging.

The dependence of viscoelastic behavior on normal loading was monitored through time sweep tests performed under normal stresses ranging from 1 to 4 kPa (**Fig.3 g-h**). For both calf and cow thymus, increasing the applied normal stress led to higher G′ and G″ values, with the most pronounced increase occurring between 1 and 2 kPa. In calf thymus, this transition corresponded to an approximately 9-fold increase in G′, followed by a more limited increase at higher normal stresses. In cow thymus, G′ also increased markedly between 1 and 2 kPa and continued to rise at higher loads, although with a less steep slope. G″ followed the same general trend, increasing strongly from 1 to 2 kPa and then remaining within a narrower range.

Across all loading conditions, cow thymus showed higher G′ and G″ values than calf thymus. The difference was particularly evident at 1 kPa, where G′ was approximately 10-fold higher in cow thymus, and remained detectable at higher normal stresses, with G′ values about 3–4-fold higher in cow than in calf thymus between 2 and 4 kPa. These data show that thymic tissue is sensitive to normal loading and that the higher stiffness of aged thymus is maintained across the entire range of applied stresses.

### 3.3 Age-associated lipid remodelling extends beyond bulk lipid accumulation

Untargeted lipidomics revealed age-associated differences in thymic lipid composition at both global and molecular levels. Overall, the relative abundance of the detected lipids was 11% higher in cow thymus compared with calf thymus (**Fig. 4a**). This global increase was not associated with a uniform variation across all lipid species. Indeed, fold-change analysis identified a total of 267 lipid species, of which 44 were significantly different between calf and cow thymus (**Fig. 4b**). These significantly altered species accounted for 16.5% of all annotated lipid species and were evenly distributed between increased and decreased lipids, with 22 species showing higher abundance and 22 species showing lower abundance in cow thymus compared with calf thymus.

**Fig. 3.**
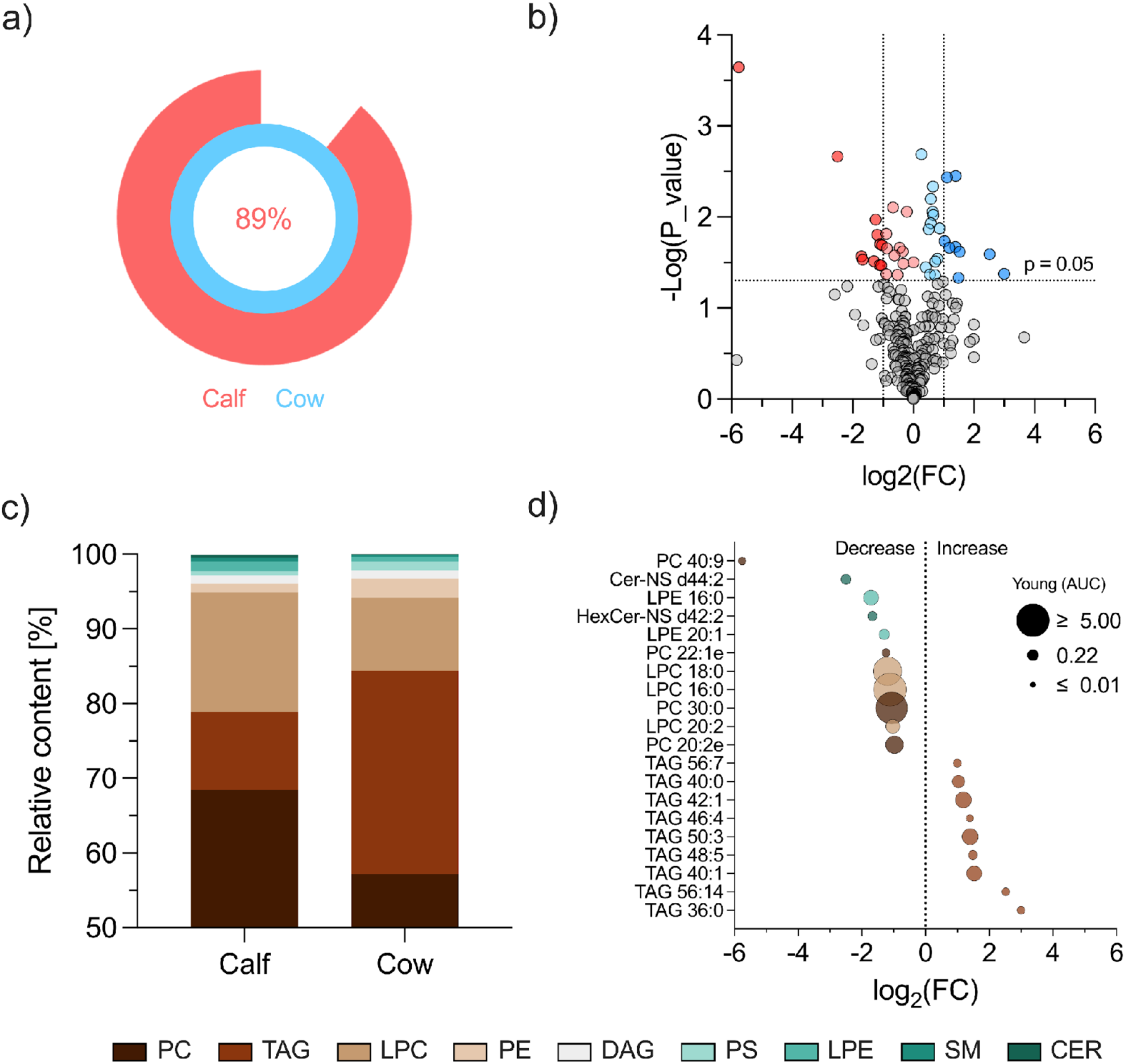
(**a**) Relative lipidic content in calf thymus normalized to cow thymus (set to 100%). (**b**) Volcano plot showing lipid variations expressed as fold change (FC) of the area under the curve (cow vs calf); Significantly decreased and increased lipids in cow relative to calf are shown in red and blue, respectively; darker shades indicate changes of at least twofold. (**c**) Distribution and relative abundance of lipid classes composing the lipid species found to be significantly different between calf and cow thymus. (**d**) Lipid species exhibiting a twofold increase or decrease (log_2(_FC) < −1 or log_2(_FC) > +1, *p < 0.05*) in AUC in cow compared to calf, with bubble size proportional to the AUC measured in calf.

The significantly altered lipids belonged to nine lipid classes: phosphatidylcholine (PC), triacylglycerol (TAG), lysophosphatidylcholine (LPC), phosphatidylethanolamine (PE), diacylglycerol (DAG), phosphatidylserine (PS), lysophosphatidylethanolamine (LPE), sphingomyelin (SM), and ceramide (Cer). Among the significantly altered species, TAGs represented the most populated class, accounting for 17 species, followed by PC with 10 species, while LPC and LPE each accounted for 4 species. Among these classes, TAGs showed the most evident increase with ageing, rising from 11% of the significantly altered lipid pool in calf thymus to 28% in cow thymus. Conversely, LPC and PC decreased in cow thymus by 39% and 16%, respectively, compared with calf thymus (**Fig. 3c**). Other lipid classes, although representing minor fractions of the total altered lipid pool, also showed marked age-associated changes. In particular, ceramides represented the least abundant class in calf thymus, accounting for 0.4% of the altered lipid pool, and decreased to 0.1% in cow thymus, corresponding to a 75% reduction. Similarly, LPE decreased from 1.3% in calf thymus to 0.7% in cow thymus, corresponding to a 46% reduction. Sphingomyelins also showed a marked decrease, declining from approximately 0.5% of the altered lipid pool in calf thymus to 0.2% in cow thymus.

When considering only lipid species showing at least a twofold increase or decrease in AUC (i.e., log₂(FC) < −1 or log₂(FC) > 1), 20 lipid species were identified (**Fig. 4d**). All lipid species increased in cow thymus belonged to the TAG class. Among them, TAG 36:0 displayed the largest relative increase, rising from an AUC of 0.044 ± 0.007 in calf thymus to 0.354 ± 0.026 in cow thymus, corresponding to an approximately eightfold increase (log₂(FC) = 2.99). In terms of absolute abundance, TAG 50:3 was the most abundant TAG among the increased species in cow thymus, with an AUC of 2.00 ± 0.72, corresponding to a 2.6-fold increase compared with calf thymus (log₂(FC) = 1.4).

By contrast, no TAG species were found among the lipids reduced in cow thymus. Decreased lipid species were instead represented by members of the PC, CER, LPC, and LPE classes. The strongest reduction was observed for PC 40:9, which decreased from an AUC of 0.041 ± 0.007 in calf thymus to values below 0.001 in cow thymus. This corresponded to an almost complete depletion of this lipid species in aged thymic tissue. Among the decreased species, LPC 16:0 and PC 30:0 showed the highest absolute abundances. LPC 16:0 decreased from 7.016 ± 1.857 in calf thymus to 3.299 ± 1.190 in cow thymus, while PC 30:0 decreased from 4.612 ± 1.590 to 2.203 ± 0.711, respectively.

### 3.4 Age-associated remodelling of elemental milieu in the thymic microenvironment

ICP–MS analysis was performed to quantify a panel of 30 metallic elements in calf and cow thymic tissues. Among these, 14 elements were detected after tissue mineralization above the limit of detection (**Tab.SI 1**). Interestingly, all elements identified in young thymic tissue were also detected in aged tissue, indicating that ageing did not result in the appearance or loss of the detectable metal species. Similarly, both calf and cow thymuses displayed a conserved elemental composition, with K, Na, Mg and Ca consistently representing the most abundant ions, whereas Ni was the least abundant detected element in both conditions.

Rank-based comparison further confirmed the overall stability of the elemental hierarchy between young and aged tissues. The four most abundant elements maintained the same ranking in calf and cow samples, while only minor rearrangements were observed among elements present at lower concentrations, particularly below 30 ng/mg tissue. These included the inversion of Zn and Fe, with Fe ranking above Zn in cow thymus, as well as the inversion of Cu and Al. Additional changes were observed among low-abundance elements, with Cr shifting to a higher relative position in cow tissue compared with calf tissue.

Despite this qualitative conservation, absolute quantification revealed a marked reduction in the total metal content of aged thymic tissue. Cow thymus contained approximately 64% of the total metal content measured in calf thymus (**Fig. 5a**). This reduction reflected a broadly distributed decrease across the elemental profile, with 11 out of 14 detected elements showing lower concentrations in cows compared with calf thymus. Conversely, only Al, Ni and Cr showed an opposite trend, increasing with age (**Fig. 5b**).

**Fig. 5.**
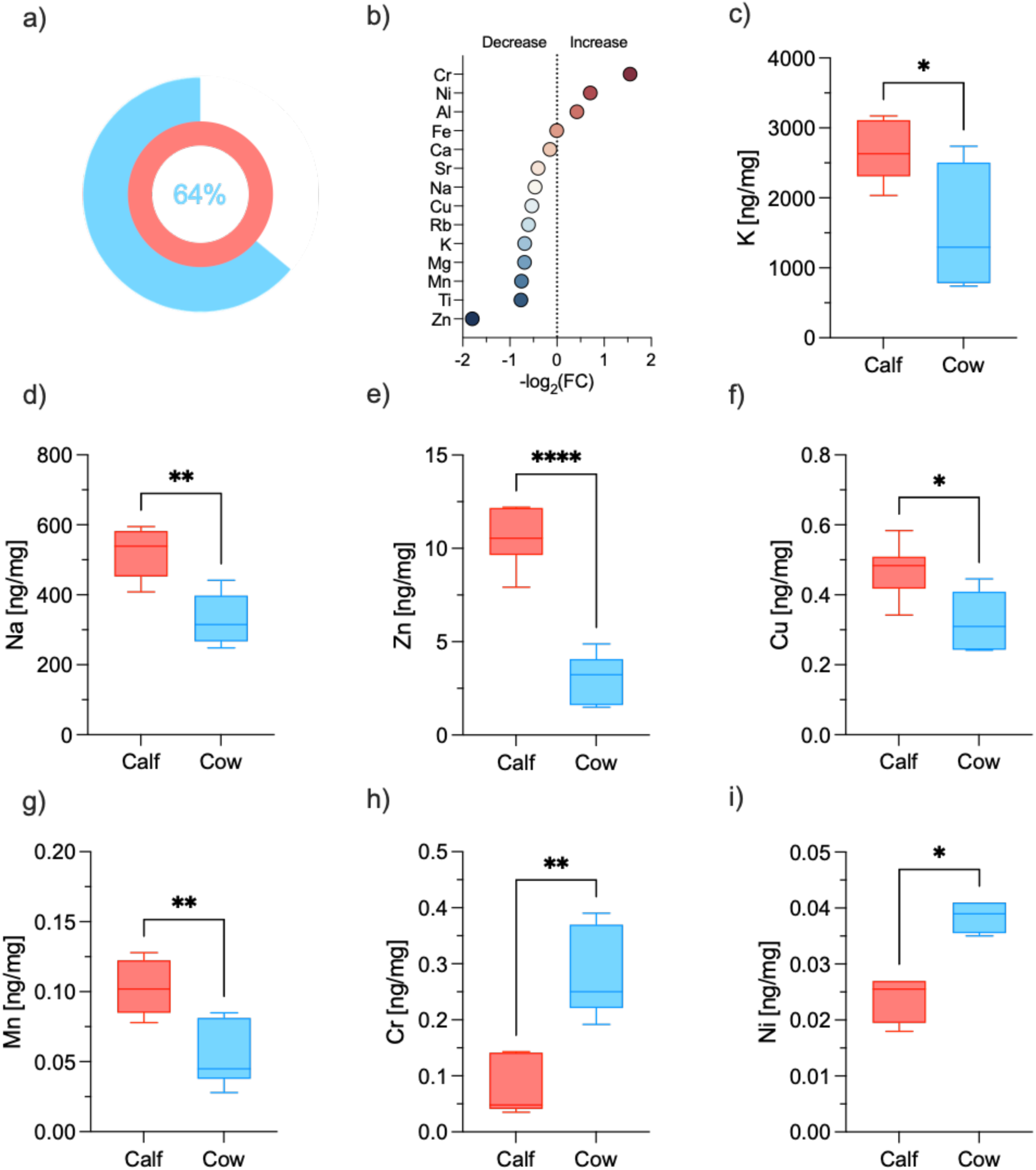
(**a**) Relative metal ion content in cow thymus normalized to calf thymus (set to 100%). A color code is used to distinguish between calf (red) and cow (blue). (**b**) log2 fold-increase of metal ion content between calf and cow thymus (**c–i**) Comparison of potassium (K), sodium (Na), magnesium (Mg), calcium (Ca), zinc (Zn), copper (Cu), and manganese (Mn), chromium (Cr) and nickel (Ni) content in calf and cow thymus.

Among the elements that decreased with age, K, Na, Zn, Cu and Mn showed the most pronounced and statistically significant reductions (**Fig. 5c–i**). In particular, Zn exhibited the strongest decrease, dropping from 11 ± 2 ng/mg tissue in calf to 3 ± 1 ng/mg in cow, corresponding to a reduction of about ∼70%. The decrease in the other significantly reduced elements was more moderate, ranging from ∼29% for Na to 41% for Mn. Among the elements that increased with age, Ni and Cr were significantly higher in cow thymus. Although Ni increased by approximately 63%, its absolute concentration remained very low, rising from 0.024 ± 0.004 to 0.039 ± 0.003 ng/mg tissue. In contrast, Cr showed a more substantial relative increase, rising from 0.076 ± 0.051 to 0.223 ± 0.081 ng/mg tissue, corresponding to an approximately threefold increase in aged thymic tissue.

### 3.5 Age-associated loss of molecular spatial organization

The metabolite signal annotated via MetaSpace was 39% higher in calf thymus (41 annotated compounds) than in cow thymus (25 annotated compounds) (**Fig. 6b**). Comparison of the annotated metabolites revealed that approximately half of the metabolites detected in calf thymus were also identified in cow thymus, with amino acids and their derivatives representing the most abundant biochemical class among the shared metabolites. The relative distribution of annotated metabolite classes revealed distinct metabolic profiles between calf and cow samples (**Fig. 6c**). Cow samples were predominantly characterized by amino acids and amino acid derivatives, which accounted for 44% of the annotated metabolites, indicating a greater representation of amino acid-related metabolism compared with calves (17%). In addition, metabolites related to choline metabolism were proportionally more abundant in cows.

**Fig. 6.**
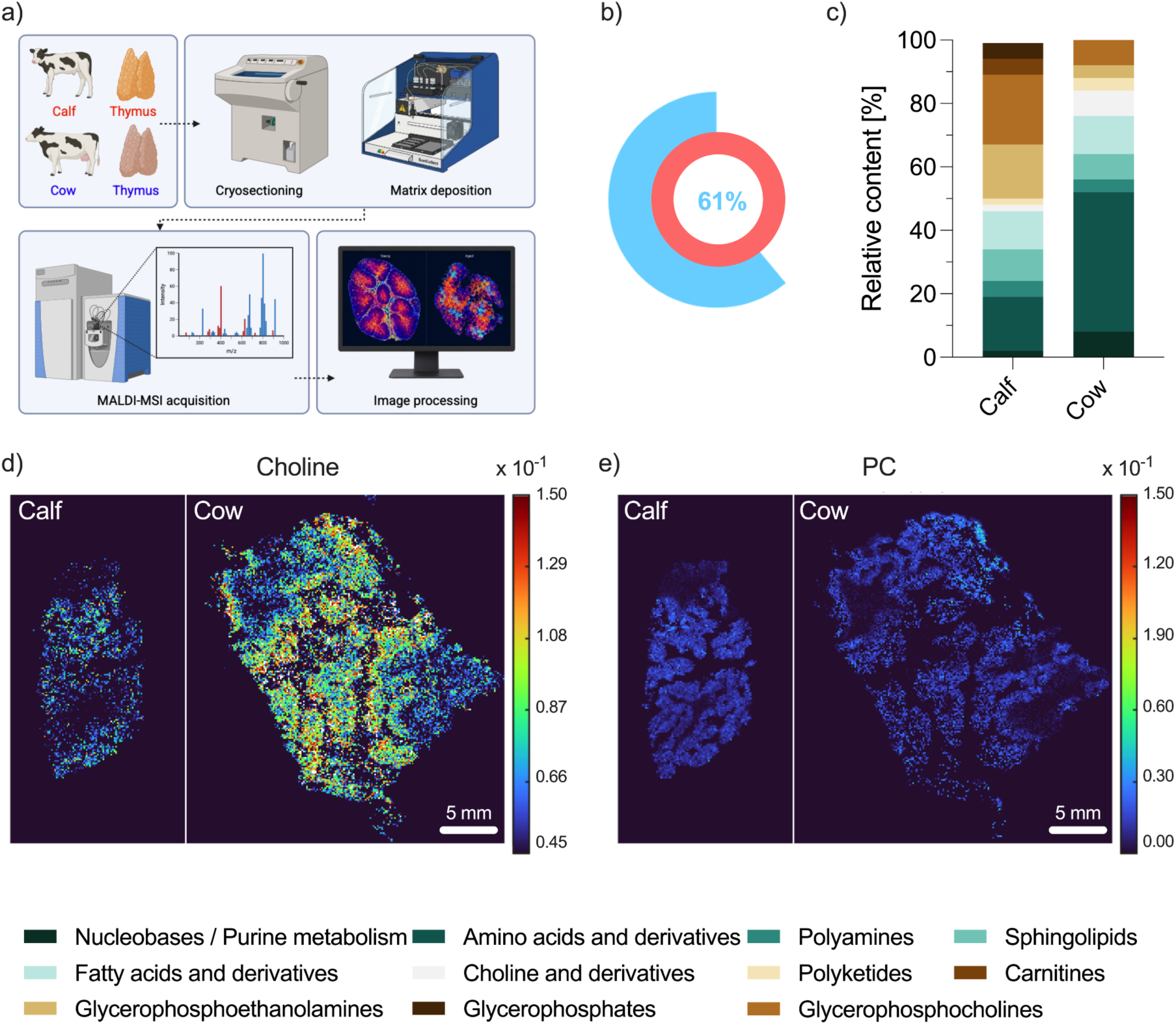
a) Schematic representation of the workflow used to acquire MALDI-MSI data. **(b)** Relative number of annotated compounds, with the number of annotations in calf samples set to 100%. **(c)** Relative distribution of annotated compounds in calf and cow thymus samples. **(d, e)** Representative ion images showing the spatial distribution of choline and phosphatidylcholine in calf and cow samples, respectively.

As a representative example, choline (*m/z* 104.1069) showed a tissue-associated signal in both groups, but with a different spatial organization: in calf thymus the signal appeared more diffusely distributed across the tissue, whereas in cow thymus it showed a more heterogeneous and patchier pattern, with areas of focal enrichment interspersed with lower-signal regions (**Fig. 6d**). In contrast, calf samples exhibited a higher proportion of membrane-associated lipids, particularly glycerophosphocholines (22% vs. 8%) and glycerophosphoethanolamines (17% vs. 4%). Consistent with this class-level difference, the representative phosphatidylcholine ion PC (isomers, *m/z* 760.5851) was detected across thymic sections in both groups, but displayed a distinct spatial arrangement, appearing relatively diffuse in calves and more discontinuous in cows, where the signal was organized in a punctate pattern with localized hotspots (**Fig. 6e**). Fatty acids and their derivatives represented an equivalent proportion of the annotated metabolites in both groups (12%), while sphingolipids showed only minor differences (10% in calves and 8% in cows). Metabolites related to purine metabolism and polyketides were proportionally more abundant in cows, whereas carnitines and glycerophosphates were detected exclusively in calves (5% each).

## Discussion

This study introduces a biomaterials-centred perspective on thymic involution by showing that ageing is accompanied by coordinated remodelling of the thymus as a physical, chemical, and spatial microenvironment[18]. The calf-to-cow comparison was used as a proof-of-concept model to determine whether age-associated thymic remodelling can be resolved as a change in tissue material state and to identify which dimensions of that state can inform biomaterials-based investigations and the engineering of thymus-inspired materials.

Because rheological measurements of native soft tissues are sensitive to storage and loading conditions[43], the effects of frozen storage, normal stress, and biological variability were examined before interpreting the age-group comparison (**Fig. 3**). Samples stored at -80 °C retained their predominantly elastic behaviour and showed no significant loss of mechanical properties for up to 3 weeks, supporting the use of frozen bovine thymic samples in comparative rheological studies when fresh tissue is unavailable. The measured moduli varied with applied normal stress, demonstrating that experimental preloading affects the apparent mechanical response. Accordingly, the lowest tested stress (1 kPa) was selected for subsequent analyses to minimize setup-induced pre-compression and obtain values more representative of the native, near-unloaded tissue. Although inter-and intra-animal variability was observed, the differences were not statistically significant, indicating that the group-level mechanical changes were primarily associated with ageing rather than sample variability.

Recent work using thymus-mimetic matrices has shown that human progenitor T-cell differentiation responds to the mechanical resistance of the engineered niche and that modulus and stress-relaxation behaviour can be controlled orthogonally[38], [44], [45]. Nevertheless, quantitative information on thymic biomechanics, particularly in relation to ageing, remains scarce. Here, ageing increased both the elastic storage modulus and viscous loss modulus by more than one order of magnitude without altering their relative balance, as tan δ remained substantially unchanged. Thus, the aged thymus retained the same broadly solid-like viscoelastic character as young tissue but exhibited markedly greater stiffness. Stiffness alone, however, did not capture the full mechanical phenotype: the linear viscoelastic region contracted in aged tissue, which entered the nonlinear regime at lower strains. This transition therefore involved two related but non-equivalent changes: greater resistance to small deformations and an earlier loss of linear mechanical behaviour as deformation increased. Comparable age-associated changes have been described in lung and arterial tissues, placing the thymic findings within a broader pattern of mechanical remodelling across ageing soft tissues[46], [47]. From a biomaterials perspective, these findings expand the relevant mechanical design space beyond stiffness alone. A scaffold that matches the G′ and G″ values of young thymus but exhibits a restricted linear viscoelastic range may still reproduce aspects of an aged, deformation-sensitive state. Incorporating viscous dissipation, the elastic-to-dissipative balance, and deformation tolerance alongside elastic modulus could therefore provide more faithful mechanical benchmarks for thymus-inspired materials.

Reproducing the mechanical properties of the young thymus would address only one component of the regenerative niche. Biomaterials can also function as chemically instructive systems: lipid composition can influence membrane turnover and mitochondrial homeostasis [48], [49], [50], whereas ion availability (and particularly Zn) has been implicated in thymocyte survival, T-cell maturation, and thymic repair [51], [52]. Defining how these compositional features change with age is therefore important not merely for describing thymic involution, but for identifying biochemical conditions that could be restored, excluded, or experimentally modulated in thymus-inspired matrices. In this context, lipidomic and elemental analyses revealed substantial differences between young and aged thymus. Lipidomics showed that thymic ageing was not characterized by uniform lipid accumulation but by selective reorganization of the lipid pool (**Fig. 4**), affecting the balance between membrane-associated functional components and storage lipids. Triacylglycerols dominated among the species that increased with age, whereas PC, LPC, LPE, SM, and Cer species decreased. A similar transition from membrane-associated lipids to storage lipids has been reported in the ageing mouse thymus. In that study, triacylglycerol accumulation was more evident in whole thymic tissue than in isolated thymocytes, suggesting expansion of lipid-rich compartments within the organ rather than an intrinsic metabolic shift in developing T cells[25]. Consistently, the TAG enrichment observed here is more likely to reflect remodelling of the thymic microenvironment than a thymocyte-specific metabolic programme. Conversely, the coordinated decrease in PC, LPC, and LPE is consistent with contraction of the functional lymphoepithelial parenchyma and loss of membrane-associated lipid pools. This interpretation is supported by evidence linking ageing to reduced phosphatidylcholine synthesis and lower PC/LPC levels, changes that may impair membrane turnover and mitochondrial integrity[48], [49], [50]. The concomitant reduction in SM and Cer extends this pattern to sphingolipids, further supporting a transition from membrane-rich functional tissue towards TAG-rich storage compartments. These age-dependent changes identify lipid metabolism as a potential target of biomaterials-based strategies for thymic regeneration rather than merely a marker of atrophy. This possibility is supported by proof-of-concept studies showing that LPE administration alleviated mitochondrial injury and improved diastolic function in aged mice[53]. More directly, growth-hormone treatment in middle-aged mice reduced thymic free cholesterol and lipid-peroxidation products while increasing thymocyte numbers and reducing inflammatory and apoptotic readouts[25]. Selective incorporation of lipids or lipid-regulating cues into biomaterials and scaffolds could therefore be explored as a means of supporting functional recovery in thymus-inspired systems.

Elemental profiling identified a second compositional layer of thymic involution (**Fig. 5**). Among the age-dependent changes in thymic metal-ion content, Zn depletion was the most prominent, with an approximately 70% reduction in aged tissue. Previous studies have linked zinc deficiency or supplementation to thymocyte apoptosis, thymic atrophy, and normalization of T-cell maturation[51], [52], while zinc sensing through GPR39 has been implicated in normal T-cell development and thymic repair after acute damage [30], [32]. This work adds a tissue-level observation to this literature by showing that total thymic Zn content differs markedly between the two age-associated states. The parallel decrease in K, Na, Cu, and Mn indicates a broader contraction of the ionic and trace-element compartment during thymic involution. Because these elements contribute to cellular fluid balance and metalloenzyme-dependent metabolism, their coordinated reduction may reflect the loss of densely cellular lymphoepithelial tissue in the aged thymus. In contrast to this general depletion, Ni and Cr increased in aged tissue. Their low abundance and sensitivity to exposure, diet, and tissue retention make them less immediately interpretable as regenerative variables than Zn[54]. The elemental profile of young thymus may therefore provide an additional design parameter for biomaterials-oriented approaches to thymic regeneration, complementing mechanical and lipidomic benchmarks. Although ICP-MS measures total tissue content rather than the bioavailability or functional state of individual metals, these findings support the investigation of ion-binding and ion-releasing matrices that control local ion availability, particularly that of Zn, and allow its effects on thymic regenerative signalling to be tested.

Histology and AP-MALDI-MSI place these mechanical and compositional changes within an architectural context. The age-associated reduction in the corticomedullary ratio observed here recapitulates a well-established feature of thymic involution (**Fig. 2**), as relative contraction of the cortical compartment has been reported in both human and murine thymus[23], [55], [56]. The corticomedullary ratio is therefore not merely a morphometric descriptor. Because the cortex and medulla provide spatially distinct microenvironments for successive stages of thymocyte development and selection, disruption of their relative balance reflects loss of the compartmental organization on which thymic function depends[57]. This interpretation is consistent with the established role of these niches in guiding thymocyte migration and maturation[58].

Within this histological background, the AP-MALDI-MSI data indicate that the age-associated loss of organization also extends to the biochemical niche (**Fig. 6**). Recent work combining MALDI-MSI and proteomics has independently demonstrated spatially resolved molecular domains in murine thymus[59], but their evolution with age has not been defined. Here, molecular classes retained in both age groups were not necessarily spatially conserved: choline shifted from a diffuse distribution in young tissue to focal enrichment within the aged parenchyma, whereas representative PC isomers became confined to discrete hotspots separated by low-signal regions. The molecular fragmentation identified by AP-MALDI-MSI therefore paralleled the loss of corticomedullary continuity observed histologically, indicating that involution remodels the topology through which biochemical cues are locally presented. The contrasting behaviour of choline and PC species is particularly informative because they are metabolically connected through membrane-phospholipid metabolism[60]. Their relative enrichment and depletion, together with their altered spatial distributions, are consistent with reorganization of the choline-phosphatidylcholine axis within the involuted tissue.

Bulk abundance and spatial presentation therefore provide complementary information: a molecule may remain detectable even when the local context in which cells encounter it changes. Spatial organization consequently emerges as a distinct dimension of thymic remodelling. For thymus-inspired biomaterials, composition should therefore be considered together with compartmentalization, gradients, and local presentation.

Taken together, the data define thymic involution as a coordinated shift in material state rather than as tissue loss alone. The young thymic niche is distinguished not only by lower viscoelastic moduli, but also by greater deformation tolerance, a membrane-associated lipid profile, a richer elemental milieu, and more continuous molecular organization. These features provide an integrated set of biomaterials-relevant benchmarks that can be reproduced individually or in combination in engineered systems to determine which aspects of the young microenvironment are necessary to support thymic repair and thymopoiesis.

## Conclusion

This work introduces a biomaterials-centred view of thymic involution, in which the physical properties, chemical composition and spatial organization of the tissue are considered together as interacting components of the native microenvironment. By defining how the young and aged thymus differ in their viscoelastic properties, lipid and elemental composition, metabolic profile and spatial molecular organization, this study moves beyond single-parameter descriptions of ageing and provides an integrated reference for the design of thymus-inspired materials. This framework now opens the possibility of identifying correlations between microenvironmental features and cellular outputs and, through their controlled manipulation in engineered models, testing individual and combined causal effects on thymic function. This could reveal which features of the young environment should be restored, preserved or reproduced to improve tissue repair and support thymopoiesis after ageing or injury.

## Authors contribution

**A. Di Bernardo,** Investigation, Data Curation, Visualization, Original Draft Writing; **C. Zanirato** Investigation, Data Curation, Visualization; **F. Briatico,** Investigation, Data Curation,Writing - Original Draft; **P. Petrini,** Investigation, Data Curation,Writing - Original Draft; **C. S. Butnarasu,** Investigation, Data Curation, Visualization,Writing - Original Draft; **F. Leo,** Investigation, Data Curation; **C. Ambrogio**, Data Curation,Visualization; **E. Patrucco**, Data Curation, Visualization; **F. Oliva,** Data Curation, Visualization, Original Draft Writing; **A. Locatelli**, Data Curation, Visualization, Original Draft Writing; **A. Passoni**, Investigation, Data Curation, Original Draft Writing; **C. Medana**, Methodology, Data Curation, Writing - Original Draft, Writing - Review & Editing; **S. Visentin**, Methodology, Data Curation, Writing - Original, Draft Writing - Review & Editing; **L. Sardelli** Conceptualisation, Methodology, Data curation, Visualization, Writing - Original Draft Writing - Review & Editing, Supervision.

## Notes

### Competing Interest Statement

The authors have declared no competing interest.

