## Supplementary Information for "An integrated biomaterials-centred approach of the ageing thymic microenvironment reveals design principles for regenerative biomaterials"

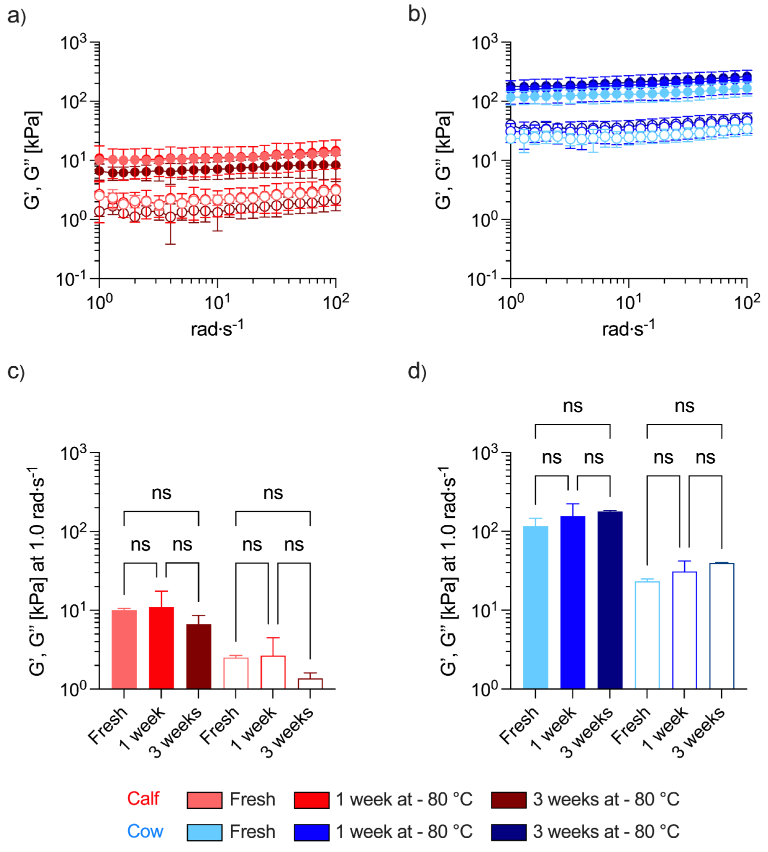


**Fig.1-SI** Storage and loss moduli of calf and cow obtained from samples stored at different conditions evaluated in the whole frequency spectra between 1 and 100 rad/s (a and b, respectively) and at the angular frequency of 10 rad/s.

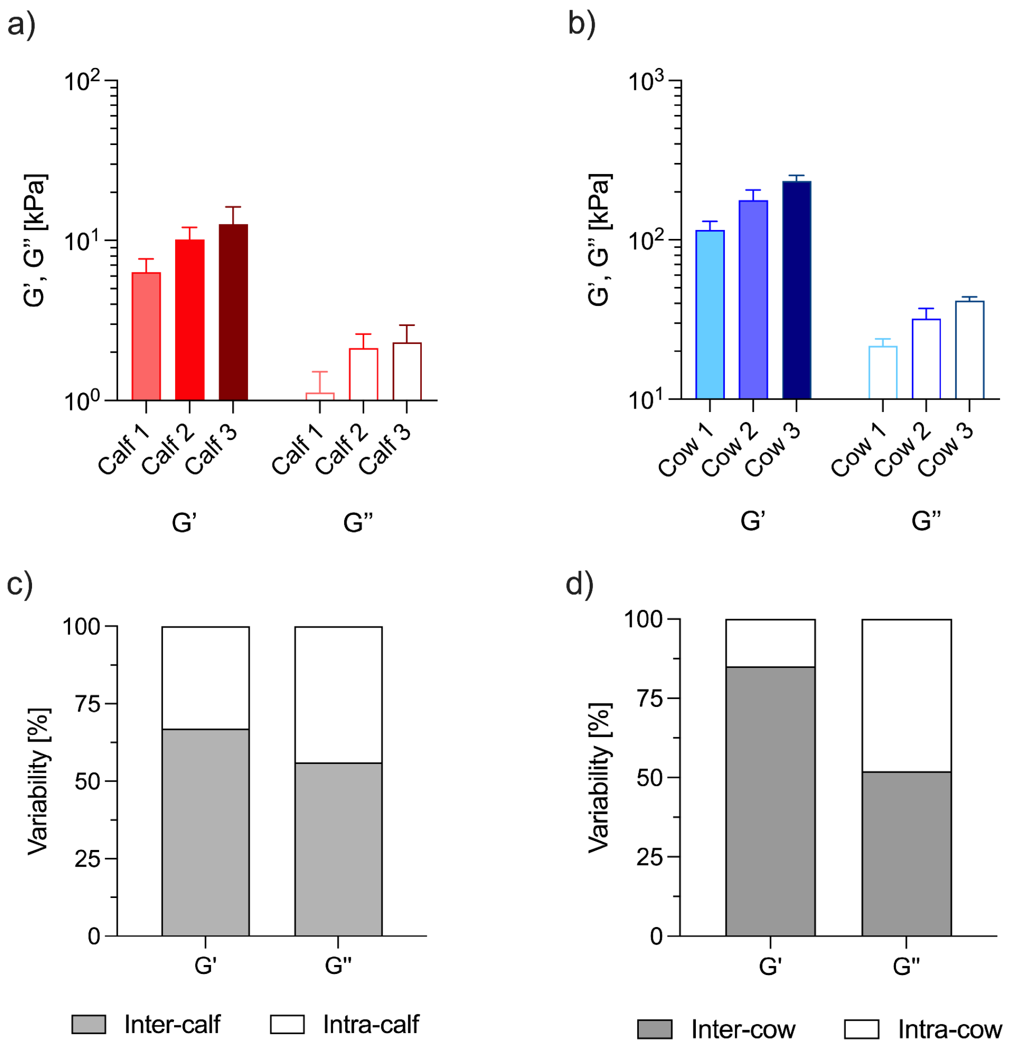


**Fig.2-SI** Storage and loss moduli of three different samples obtained from three different calves and cows and at the angular frequency of 10 rad/s (a and b) and inter and intra variability coefficients (c for calf and d for cow, respectively)

**TableSI 1**. Metal ions content in calf and cow thymus quantified by inductively coupled plasma-mass spectrometry

| | Calf | | | Cow | | |
|---|---|---|---|---|---|---|
| **ng/mg** | **average** | **SD** | **n** | **average** | **SD** | **n** |
| **K** | 2493,044 | 741,766 | 6 | 1548,308 | 791,883 | 6 |
| **Na** | 454,364 | 145,422 | 6 | 329,261 | 69,199 | 6 |
| **Mg** | 124,442 | 50,796 | 6 | 77,199 | 50,313 | 6 |
| **Ca** | 32,294 | 4,098 | 6 | 29,105 | 5,980 | 6 |
| **Zn** | 8,930 | 3,362 | 6 | 5,047 | 3,284 | 6 |
| **Fe** | 8,207 | 1,433 | 6 | 8,169 | 1,601 | 6 |
| **Cu** | 0,470 | 0,076 | 6 | 0,325 | 0,078 | 6 |
| **Mn** | 0,091 | 0,033 | 6 | 0,054 | 0,022 | 6 |
| **Rb** | 3,154 | 0,967 | 6 | 2,075 | 1,063 | 6 |
| **Al** | 0,291 | 0,302 | 6 | 0,406 | 0,408 | 6 |
| **Ti** | 0,134 | 0,051 | 6 | 0,080 | 0,048 | 6 |
| **Sr** | 0,060 | 0,018 | 6 | 0,045 | 0,008 | 6 |
| **Ni** | 0,025 | 0,003 | 6 | 0,039 | 0,003 | 6 |
| **Cr** | 0,041 | 0,001 | 6 | 0,019 | 0,000 | 6 |
| **Pb** | < LOD | < LOD | 6 | 0,018 | 0,001 | 6 |
| **V** | ND | ND | 6 | ND | ND | 6 |
| **Co** | ND | ND | 6 | ND | ND | 6 |
| **As** | ND | ND | 6 | ND | ND | 6 |
| **Se** | ND | ND | 6 | ND | ND | 6 |
| **Sr** | ND | ND | 6 | ND | ND | 6 |
| **Zr** | ND | ND | 6 | ND | ND | 6 |
| **Nb** | ND | ND | 6 | ND | ND | 6 |
| **Mo** | ND | ND | 6 | ND | ND | 6 |
| **Ru** | ND | ND | 6 | ND | ND | 6 |
| **Rh** | ND | ND | 6 | ND | ND | 6 |
| **Pd** | ND | ND | 6 | ND | ND | 6 |
| **Ag** | ND | ND | 6 | ND | ND | 6 |
| **Cd** | ND | ND | 6 | ND | ND | 6 |
| **Sn** | ND | ND | 6 | ND | ND | 6 |
| **Sb** | ND | ND | 6 | ND | ND | 6 |
| **Ba** | ND | ND | 6 | ND | ND | 6 |
| **W** | ND | ND | 6 | ND | ND | 6 |
| **Pt** | ND | ND | 6 | ND | ND | 6 |
| **Hg** | ND | ND | 6 | ND | ND | 6 |
| **Tl** | ND | ND | 6 | ND | ND | 6 |
| **Bi** | ND | ND | 6 | ND | ND | 6 |
| **Th** | ND | ND | 6 | ND | ND | 6 |
| **U** | ND | ND | 6 | ND | ND | 6 |

ND = not detected
LOD = limit of detection

| m/z | Compound | Ion images |
|---|---|---|

| | | |
|---|---|---|
| 104.1069 | Choline | 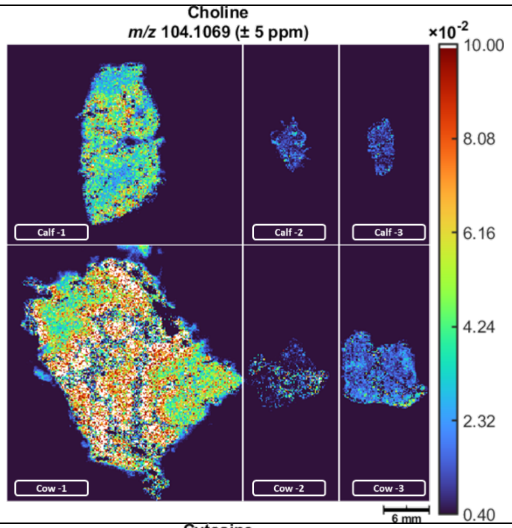 |
| 112.0505 | Cytosine | 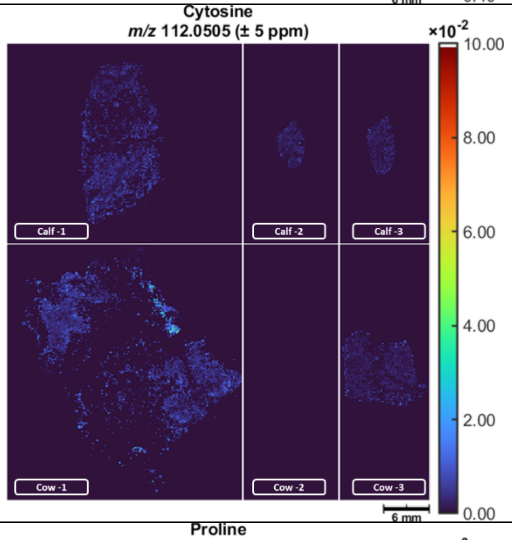 |
| 116.0704 | Proline | 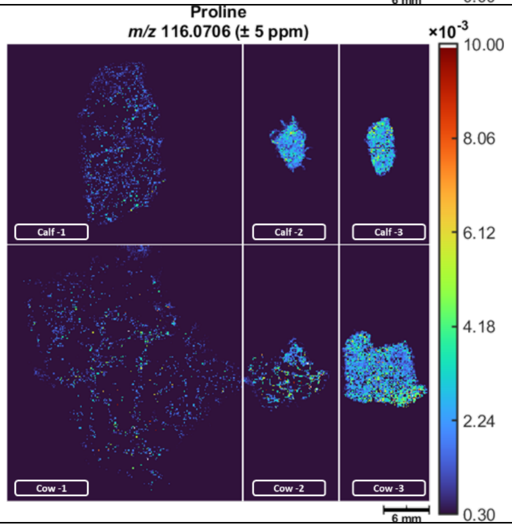 |

| | | |
|---|---|---|
| 132.1019 | Leucine/ Isoleucine | 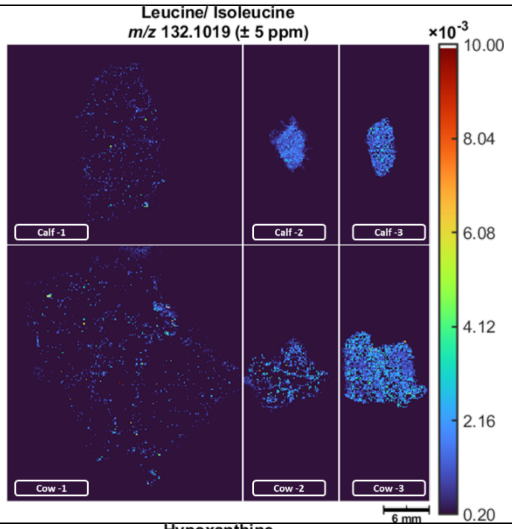 |
| 137.0458 | Hypoxanthine | 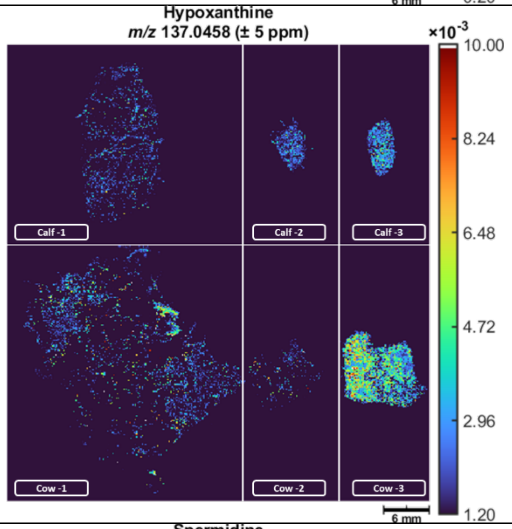 |
| 146.1652 | Spermidine | 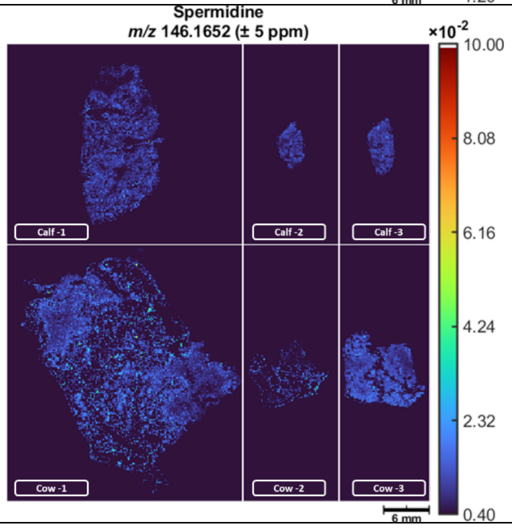 |

| | | |
|---|---|---|
| 148.0604 | Glutamic acid/<br>Acetylserine | 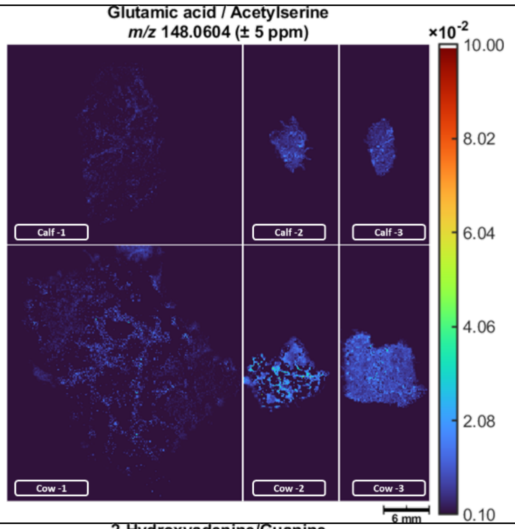 |
| 152.0567 | 2-Hydroxyadenine/<br>Guanine | 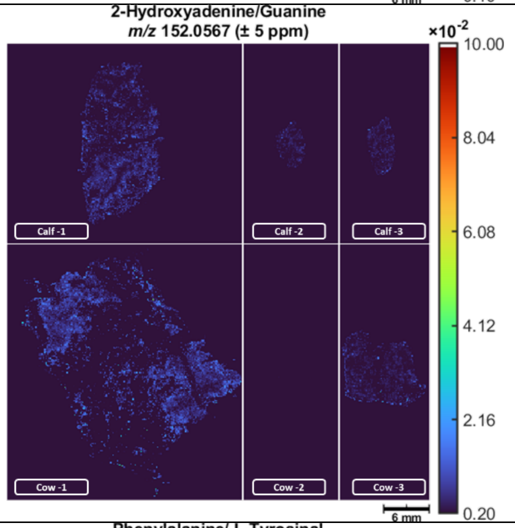 |
| 166.0862 | Phenylalanine/<br>L-Tyrosinal | 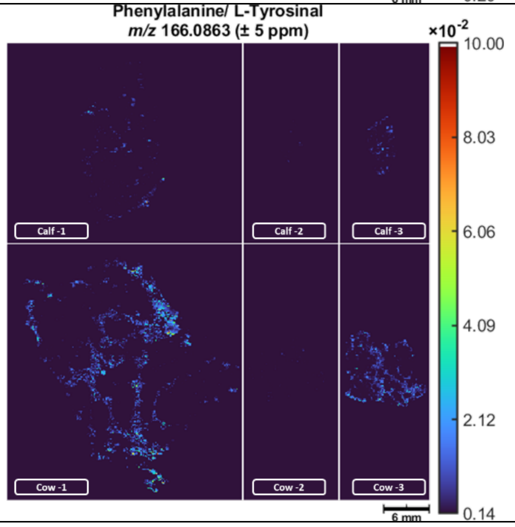 |

| | | |
|---|---|---|
| 180.0655 | Hippuric acid | 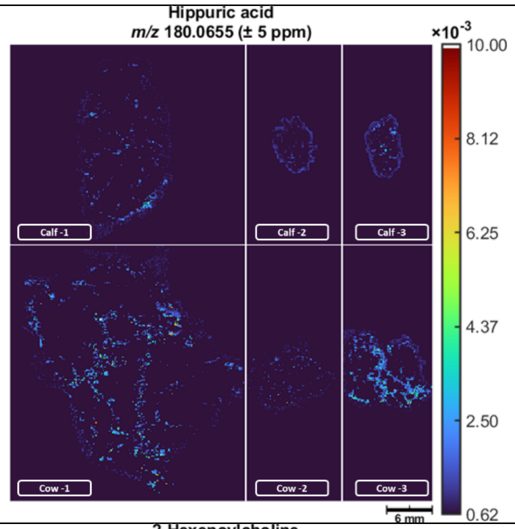 |
| 200.1645 | 2-Heneoylcholine | 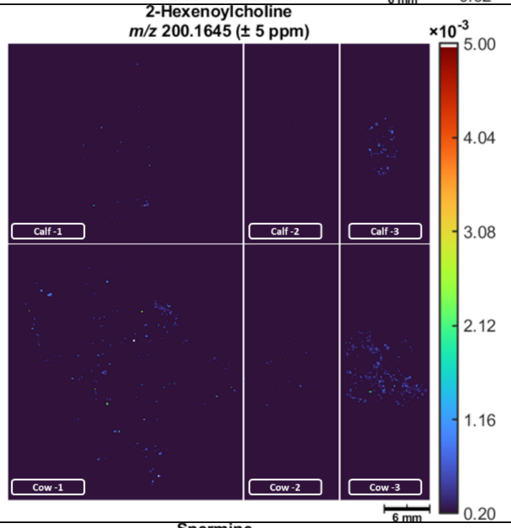 |
| 203.2230 | Spermine | 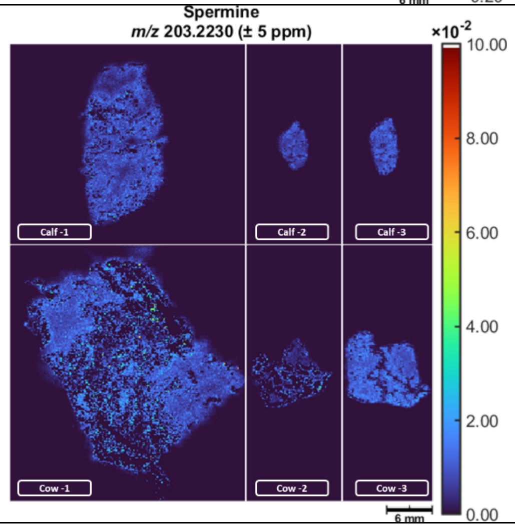 |

| | | |
|---|---|---|
| 208.0604 | 4-(2-Aminophenyl)-2,4-dioxobutanoate/ 2-Formaminobenzoylacetate | 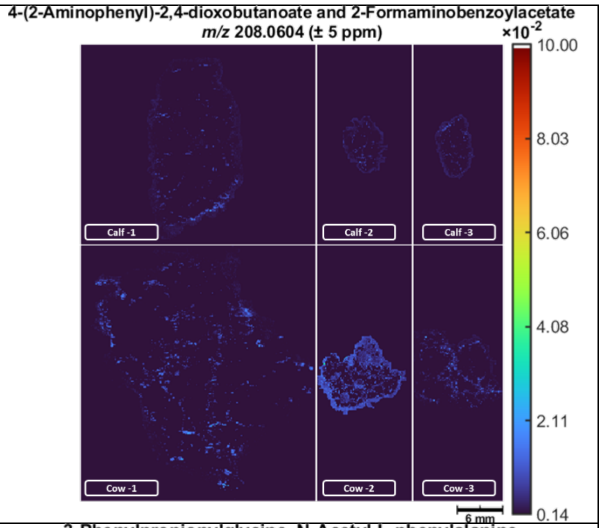 |
| 208.0968 | 3-Phenylpropionylglycine, N-Acetyl-L-phenylalanine | 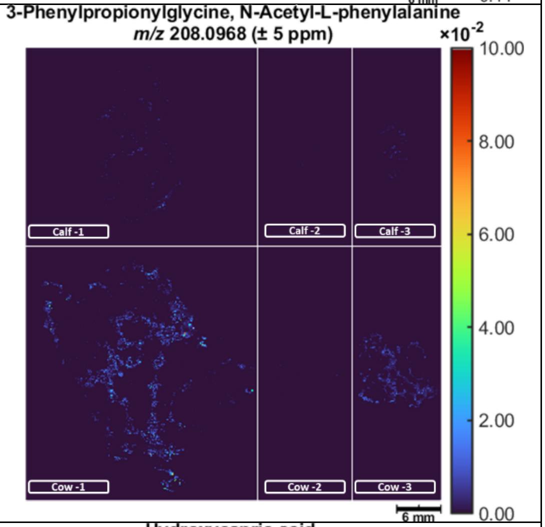 |
| 211.1304 | Hydroxycapric acid | 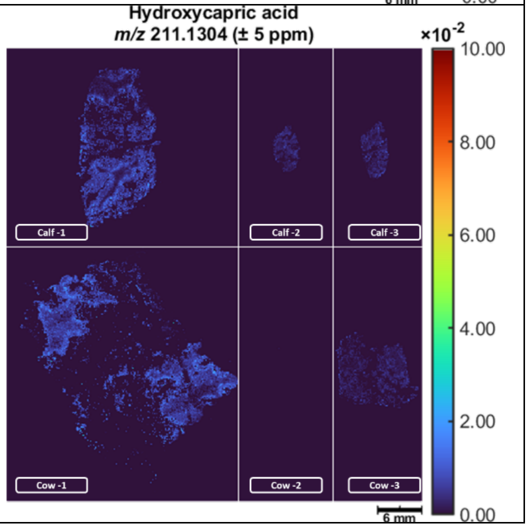 |

| | | |
|---|---|---|
| 249.1234 | 6-Hydroxymelatonin | 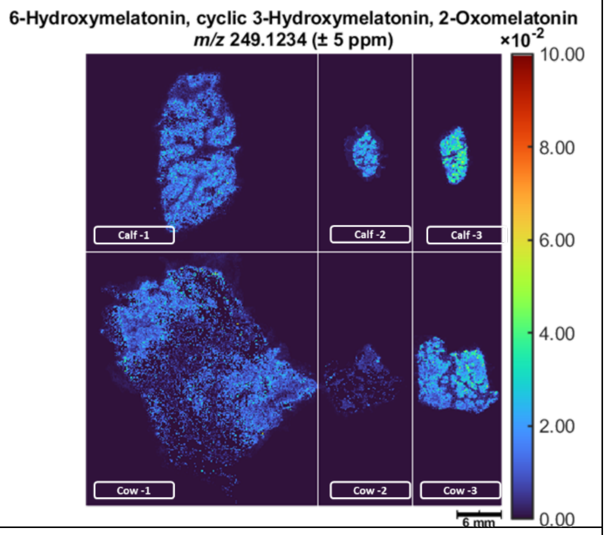 |
| 252.0866 | 4-Hydroxy-3-methoxy-cinnamoylglycine | 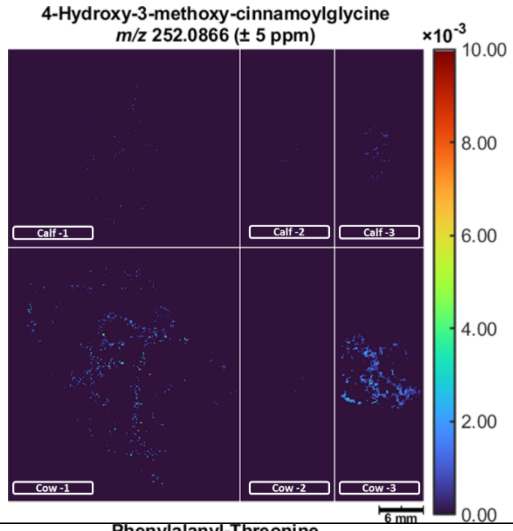 |
| 267.1339 | Phenylalanyl/Threonine | 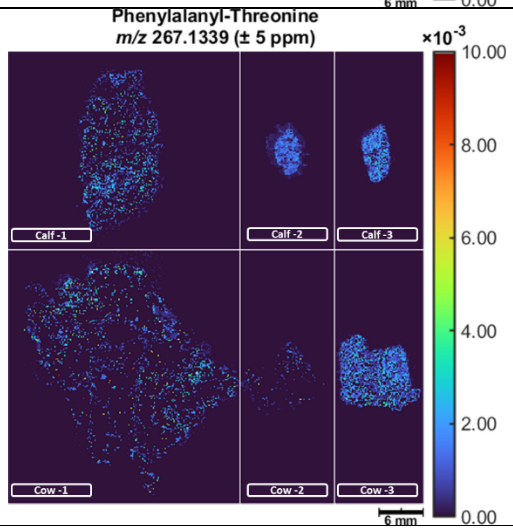 |

| | | |
|---|---|---|
| 282.2791 | Elaidamide/Oleamide | 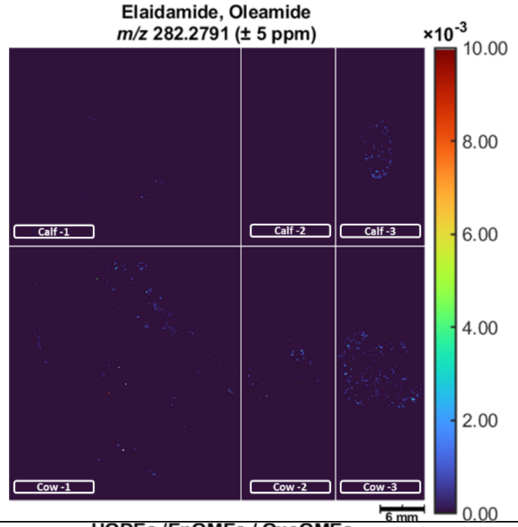 |
| 297.2424 | HODEs (Hydroxyoctadecadienoic acids) / EpOMEs (Epoxyoctadecamonoenoic acids) OxoOMEs / keto-octadecenoic acids (isomers) | 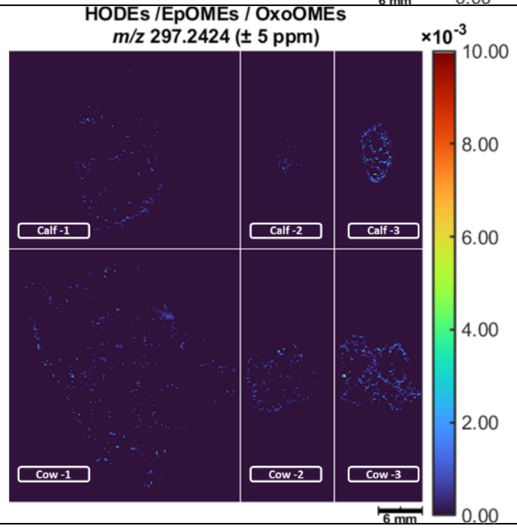 |
| 314.2690 | Palmitoylglycine | 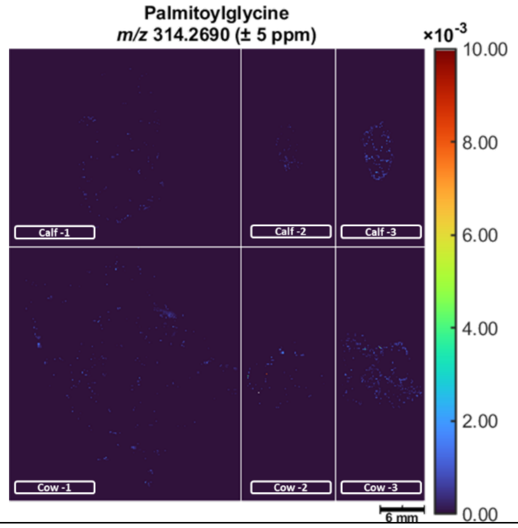 |

<table>
<tr><td>316.2846</td><td>4-hydroxy-8cis-sphingenine/ Dehydrophytosphingosine/ 6-hydroxysphingosine</td><td>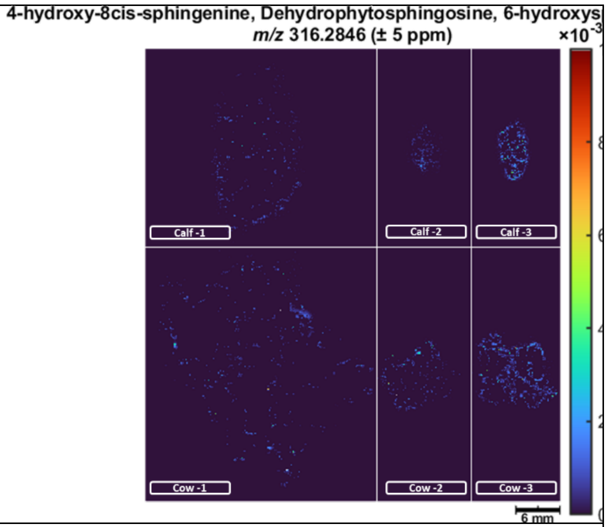
</td></tr>
<tr><td>318.3003</td><td>Phytosphingosine</td><td>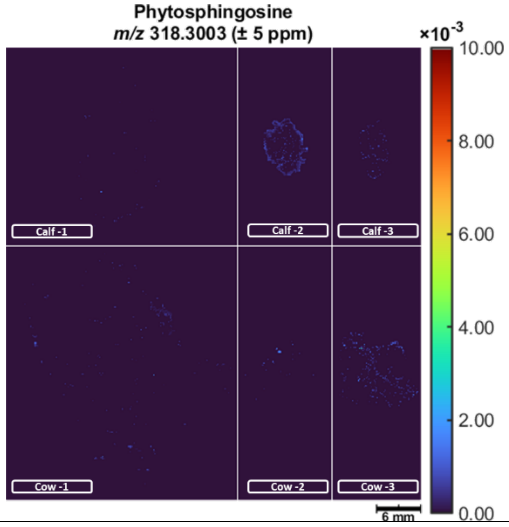
</td></tr>
<tr><td>328.2846</td><td>N-palmitoyl alanine / Margaroylglycine</td><td>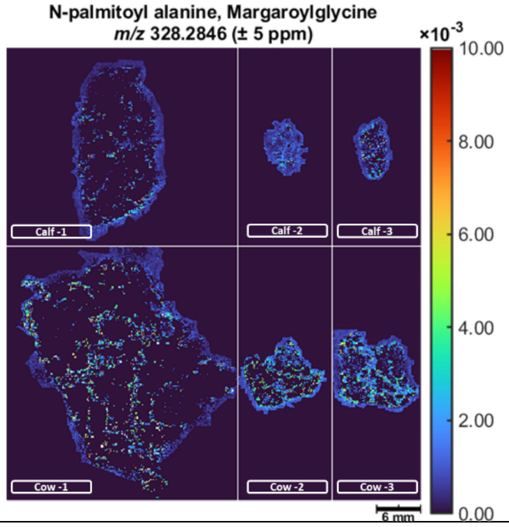
</td></tr>
</table>

| | | |
|---|---|---|
| 330.2639 | Undecanoylcarnitine / 4,8 Dimethylnonanoyl carnitine | 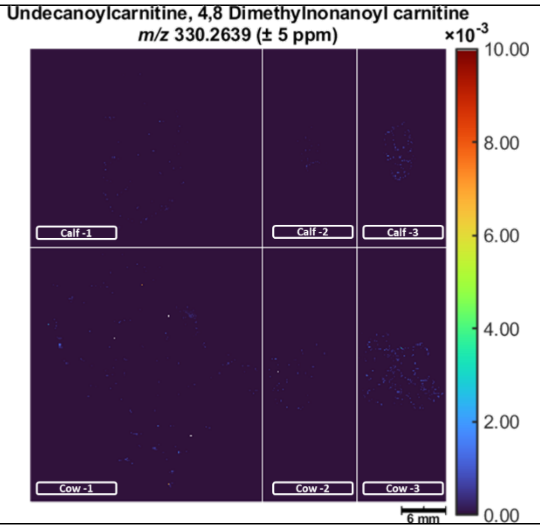  |
| 352.1656 | Tryptophyl-Phenylalanine/ Phenylalanyl-Tryptophan | 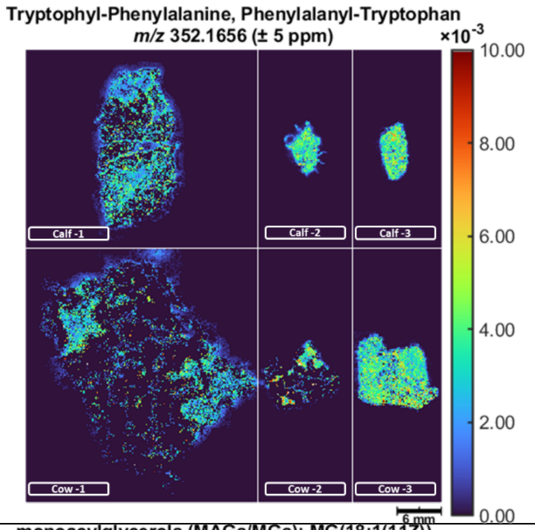  |
| 357.2999 | Monoacylglycerols (MAGs/MGs): MG(18:1(11Z)) | 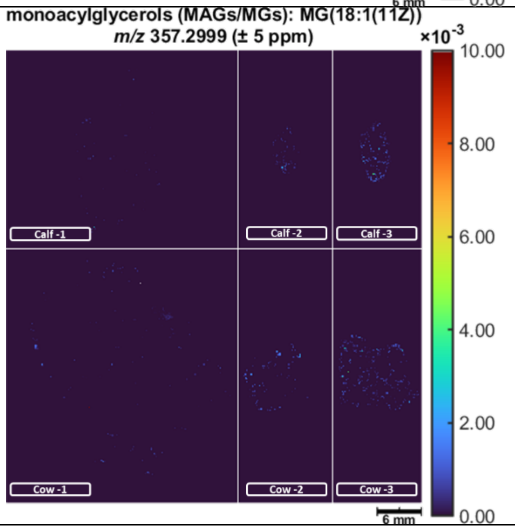  |

| | | |
|---|---|---|
| 372.3108 | N-stearoyl serine/ Tetradecanoylcarnitine | 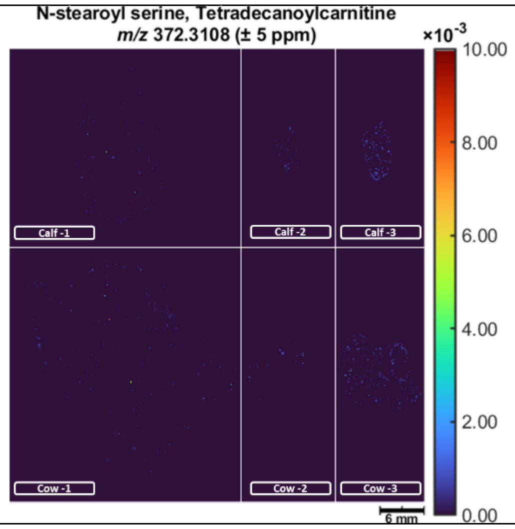 |
| 395.2557 | Cervonoyl ethanolamide | 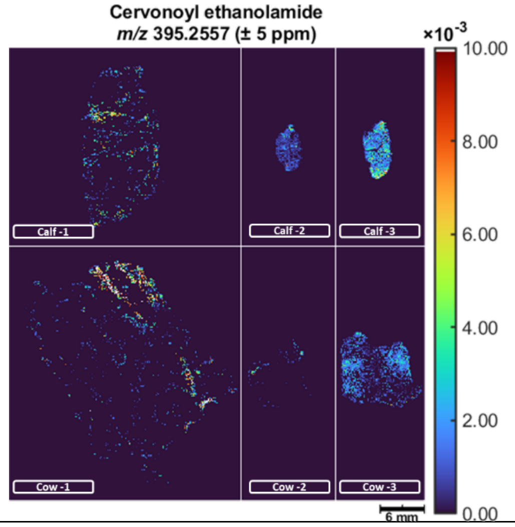 |
| 455.0036 | 4-Acetamido-4'-isothiocyanostilbene-2,2'-disulphonic acid, SITS | 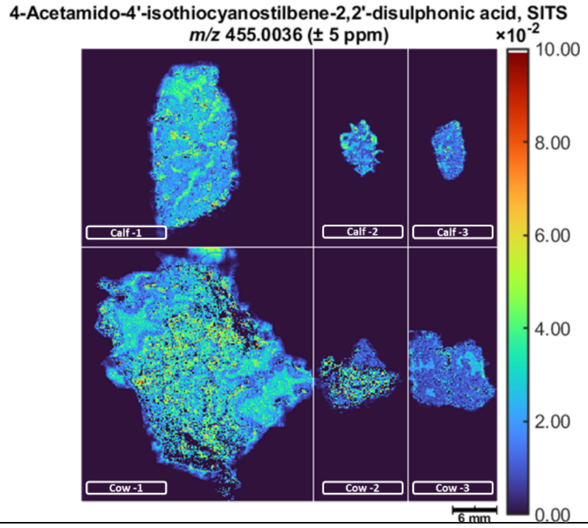 |

| | | |
|---|---|---|
| 480.3085 | PE(18:1(9Z)/0:0) | 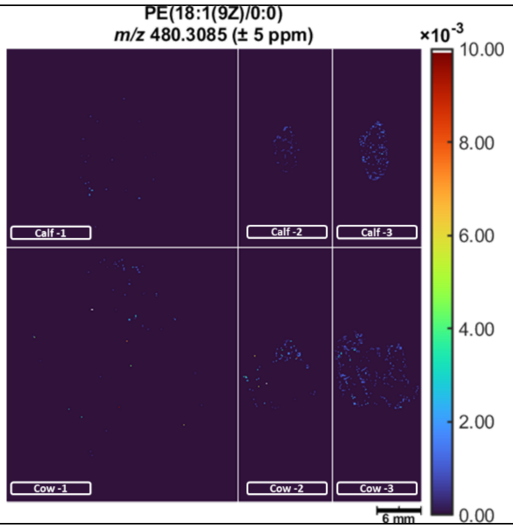 |
| 482.3241 | LysoPE(18:0/0:0) | 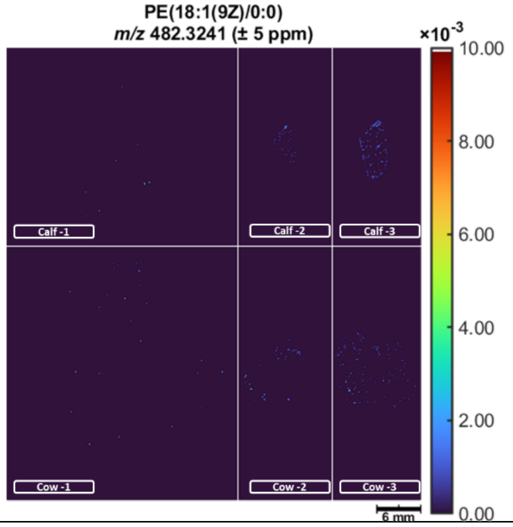 |
| 496.3398 | PC(0:0/16:0) | 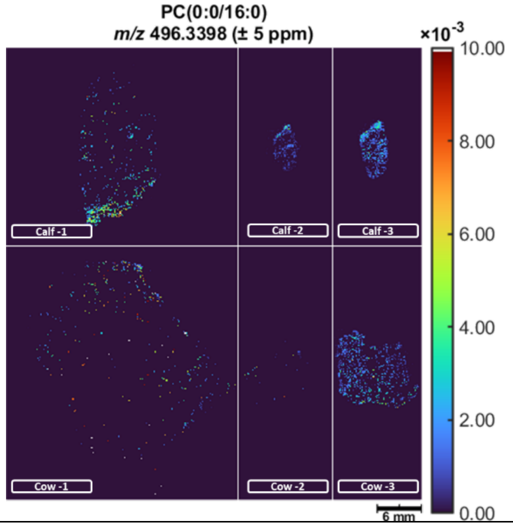 |

| | | |
|---|---|---|
| 502.2928 | LysoPE(0:0/20:4) | 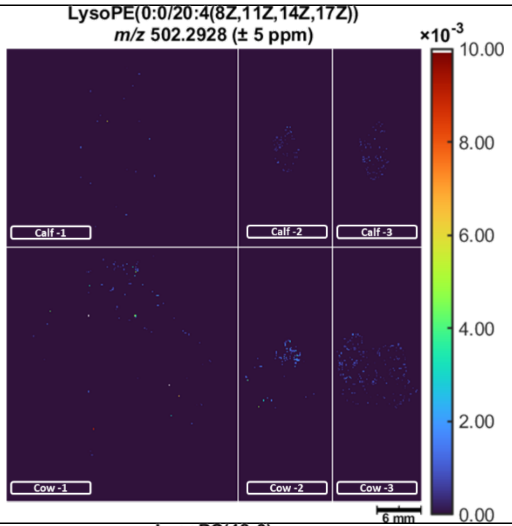 |
| 524.3711 | LysoPC(18:0) | 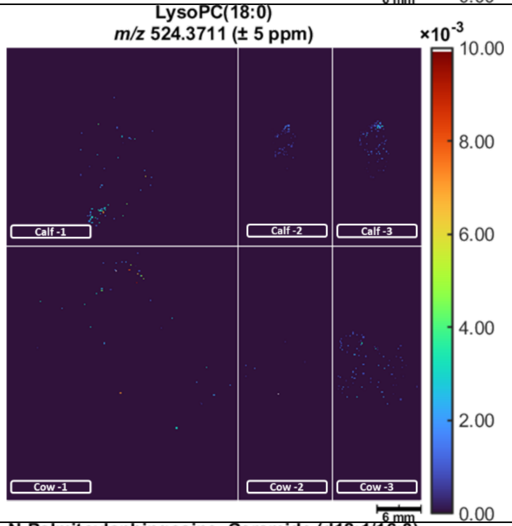 |
| 538.5194 | N-Palmitoylsphingosine / Ceramide (d18:1/16:0) | 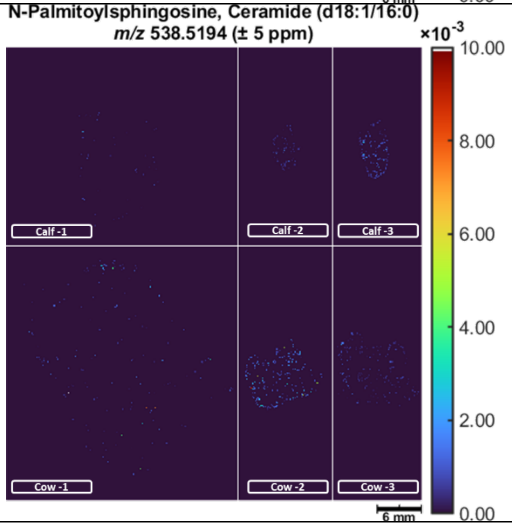 |

| | | |
|---|---|---|
| 633.4854 | PA(O-18:0/14:1(9Z)) | 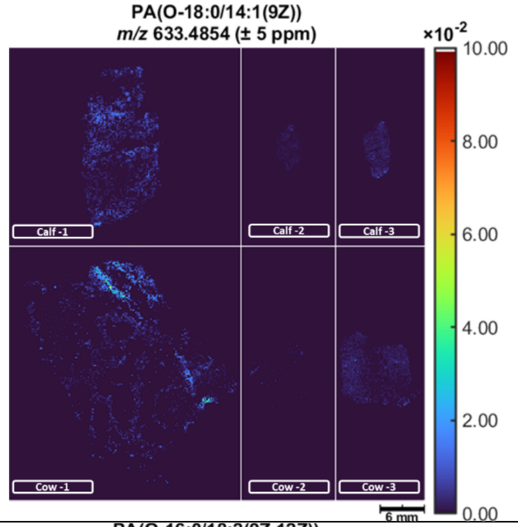 |
| 659.5010 | PA(O-16:0/18:2(9Z,12Z)) | 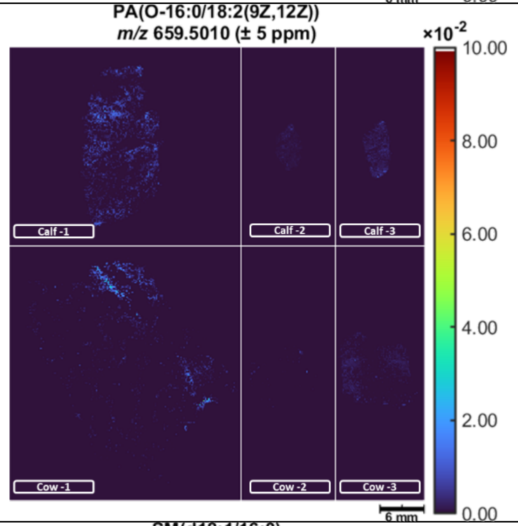 |
| 703.5749 | SM(d18:1/16:0) | 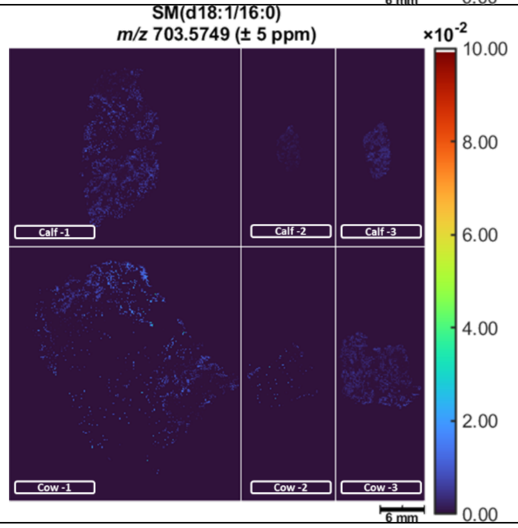 |

| | | |
|---|---|---|
| 718.5381 | PE(14:1(9Z)/20:0) | 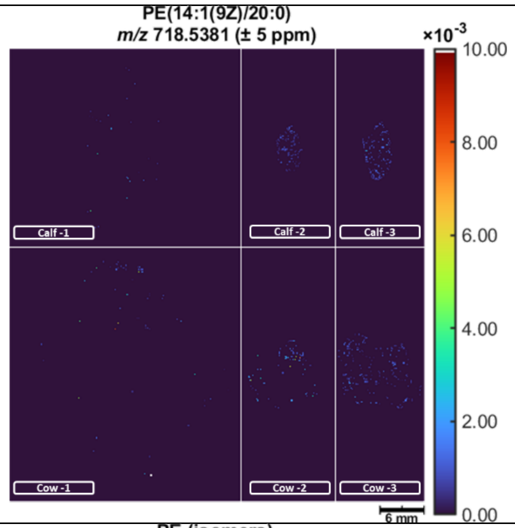 |
| 720.5538 | PE (isomers) | 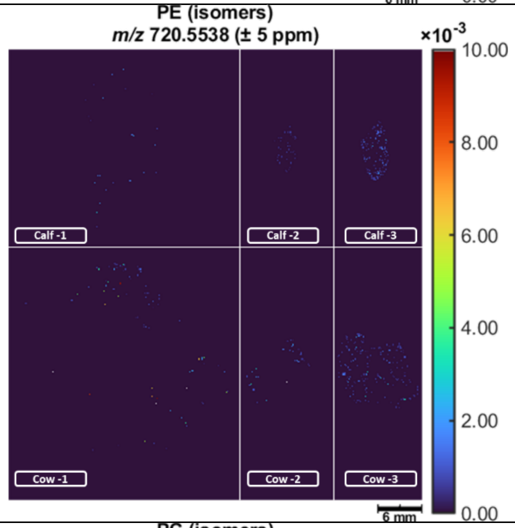 |
| 732.5538 | PC (isomers) | 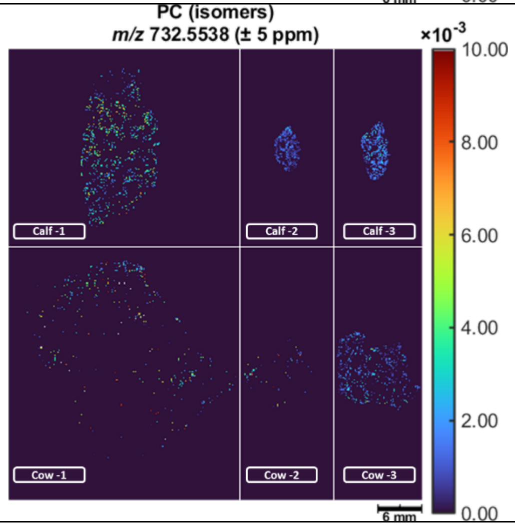 |

<table>
<tr><td>746.6058</td><td>PC(o-16:1(9Z)/18:0)</td><td>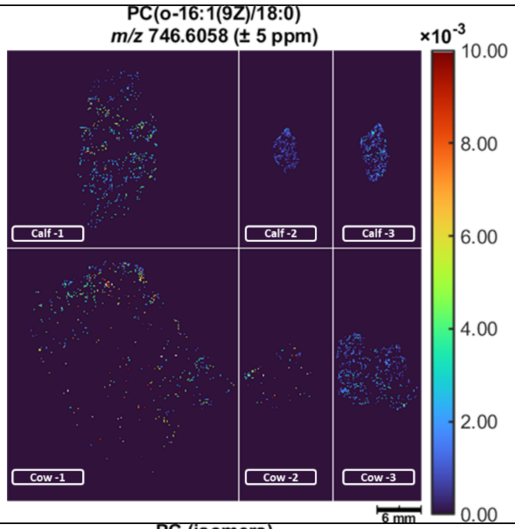
</td></tr>
<tr><td>760.5851</td><td>PC (isomers)</td><td>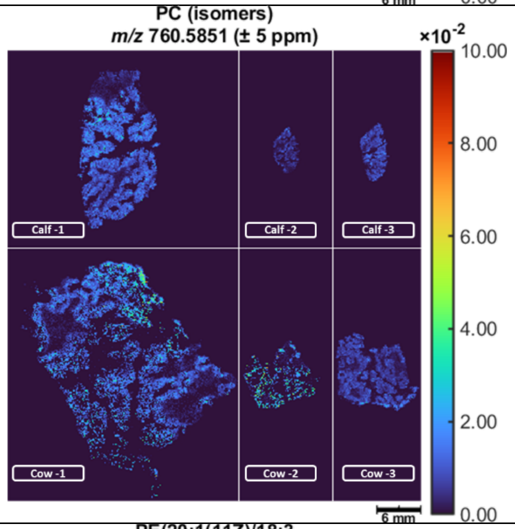
</td></tr>
<tr><td>768.5538</td><td>PE(20:1(11Z)/18:3</td><td>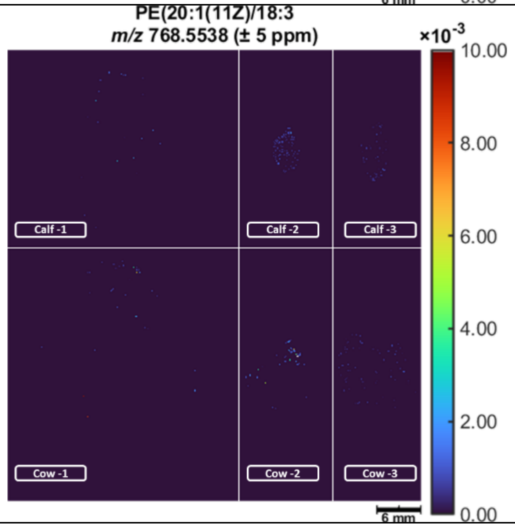
</td></tr>
</table>

| | | |
|---|---|---|
| 774.6007 | PE (isomers) | 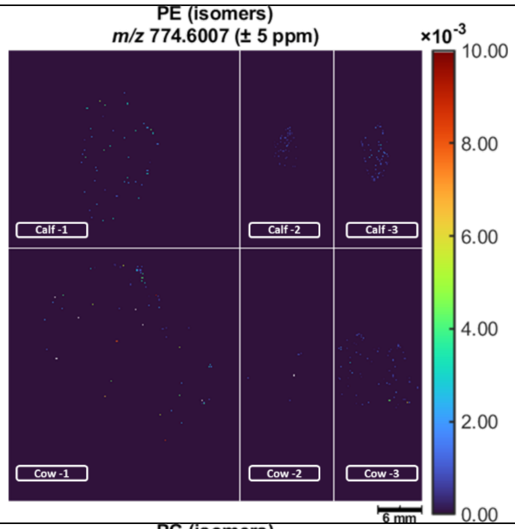 |
| 784.5851 | PC (isomers) | 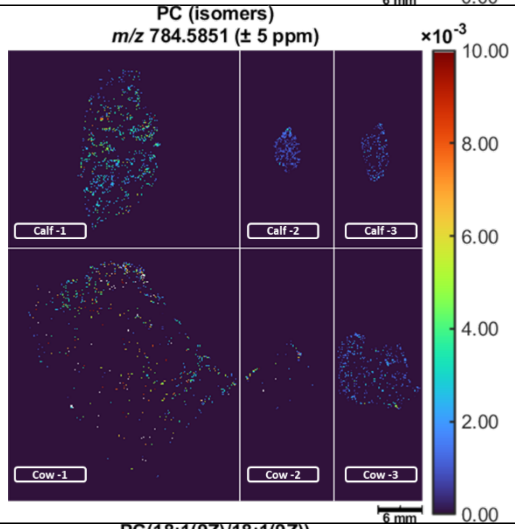 |
| 786.6007 | PC(18:1(9Z)/18:1(9Z)) | 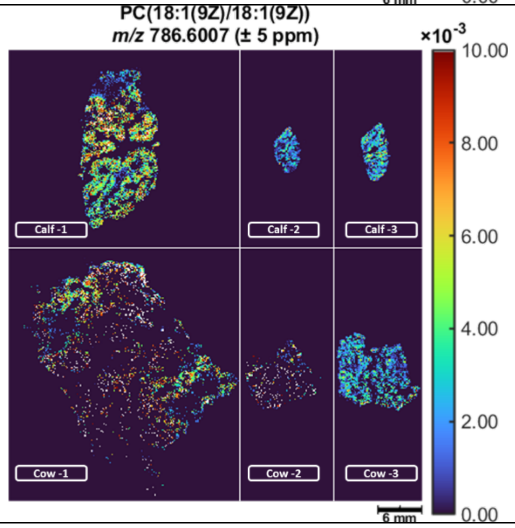 |

| | | |
|---|---|---|
| 788.6164 | PC(14:1(9Z)/22:0) | 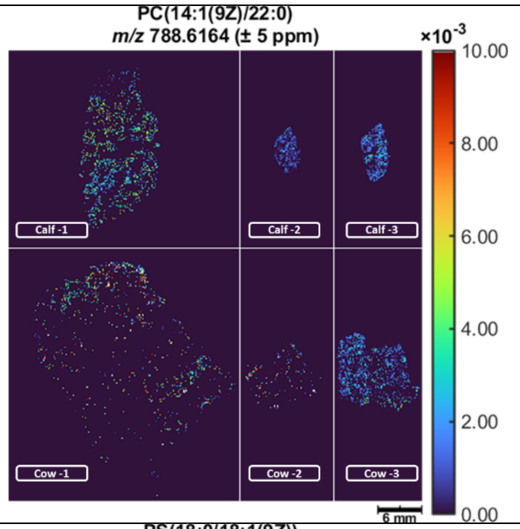 |
| 790.5593 | PS(18:0/18:1(9Z)) | 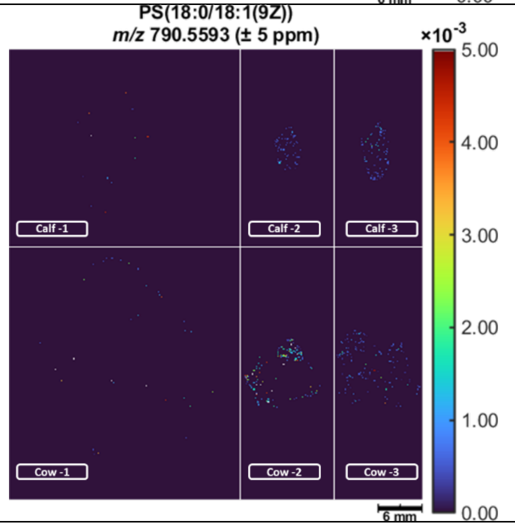 |

**Table S1.** Metabolites identified and their relative spatial distribution by MALDI mass spectrometry imaging (MALDI-MSI) in calf thymus (top) and cow thymus (bottom). For each experimental group, the first image (left) was acquired at a spatial resolution of 50 µm, whereas the other two images were acquired at 150 µm. Ion intensities were normalized to the total ion current (TIC) and are displayed using a color-coded scale (arbitrary units, a.u.); the maximum intensity value is indicated for each panel. Scale bar = 5 mm.
